# Genetic activation of ERK2 recapitulates core neurodevelopmental features of Rasopathy syndromes in mice

**DOI:** 10.64898/2025.12.15.694380

**Authors:** Kassidy E. Grover, Zoe R. Cappel, Avery Volz, Evelin M. Cotella, Kelly Smallwood, Christine A. Berryhill, Kimaya R. Raje, Austen A. Fisher, Mary Claire T. Casper, Diana Nardini, Tilat A. Rizvi, Rosa M. Salazar, Ashley Wooten, Michael T. Williams, Charles V. Vorhees, Lindsey E. Romick, Kenneth D. Greis, Yueh-Chiang Hu, Linde A. Miles, Steven P. Angus, Nancy Ratner, Carlos E. Prada, K. Nicole Weaver, Ronald R. Waclaw, J. Elliott Robinson

## Abstract

Germline pathogenic variants that activate the Ras/mitogen-activated protein kinase (MAPK) pathway cause neurodevelopmental disorders called ‘Rasopathies’. Because many affected proteins directly regulate Ras, causative mutations may alter other Ras-dependent pathways in addition to MAPK signaling. To better understand which Rasopathy sequelae result from hyperactivation of downstream MAP kinases, we engineered mice with a gain-of-function mutation in the terminal MAP kinase gene *Mapk1*, which encodes ERK2 and is associated with the recently described genetic syndrome *MAPK1*-related Rasopathy (MRR). *Mapk1* mutant mice successfully modeled key aspects of the human MRR phenotype, including small stature, facial dysmorphism, and impaired cognitive function. Importantly, they recapitulated phenotypes identified in Rasopathy models with upstream Ras activation, such as neurofibromatosis type 1 (NF1): oligodendrocyte lineage defects, reactive astrogliosis, memory deficits, and hypersensitivity to sensory stimuli. These findings emphasize the importance of downstream MAPK signaling in the pathophysiology of neurocognitive symptoms observed in Rasopathy syndromes.

## INTRODUCTION

The Ras/mitogen-activated protein kinase (MAPK) pathway is a biochemical signaling cascade that regulates several basic cellular processes, including growth, differentiation, and responses to stress^1–5^. In this pathway, extracellular ligands (e.g. hormones, cytokines, growth factors) alter cell states by binding cell surface receptors, which activate the Ras family of small GTPases by promoting the exchange of guanosine disphosphate (GDP) for guanosine triphosphate (GTP). Once active, GTP-bound Ras initiates the sequential phosphorylation of a series of serine/threonine protein kinases (Raf, MEK, and ERK). ERK (extracellular signal-regulated kinase) is the terminal MAP kinase and influences cellular function by directly phosphorylating cytosolic targets or traveling to the nucleus where it can regulate gene expression^3,4^. While somatic mutations activating the MAPK pathway are prevalent in cancers, germline variants are associated with a family of developmental disorders called ‘Rasopathies’^6^. Rasopathies include neurofibromatosis type 1 (NF1), Noonan syndrome (NS), Noonan syndrome with multiple lentigines (NSML), Costello syndrome (CS), and cardofaciocutaneous syndrome (CFC). These Rasopathy syndromes have unique and shared clinical features, such as developmental delay, craniofacial abnormalities, congenital heart defects, cognitive impairment, and an increased risk of malignancy^6–8^.

Noonan syndrome (NS) is the most common Rasopathy and has an incidence of approximately 1 in 1000-2500 live births^9^. In addition to manifestations listed above, NS patients may experience short stature, autism spectrum disorder (ASD), attention-deficit/hyperactivity disorder (ADHD)^10–13^, gastrointestinal problems^14^, etc. The genetic etiology of NS is heterogeneous and most commonly results from mutations in *PTPN11, SOS1, KRAS,* and *RIT1* genes^9^. Of these, *PTPN11* mutations are most prevalent, occurring in approximately 50% of cases^15,16^. Like NS patients, mice with gain-of-function mutations in *Ptpn11*, *Kras*, and *Rit1* genes experience cardiac defects, small size, craniofacial and hematological abnormalities, and cognitive dysfunction^17–20^. Because these mutations affect signaling upstream or at the level of the Ras GTPase, phenotypes may be caused by aberrant activation of downstream MAP kinases or other Ras-regulated proteins, such as phosphoinositide 3-kinase, Rac1, Ral, etc.^11,21,22^. Therefore, the development of transgenic mice with gain-of-function mutations in terminal MAP kinases would clarify the role of this signaling pathway in the pathophysiology of Rasopathy syndromes.

Recently, an autosomal dominant neurodevelopmental syndrome on the Rasopathy clinical spectrum with NS-like features was described in seven patients with *de novo* missense mutations in the *MAPK1* gene^23^, which encodes the terminal MAP kinase ERK2. These patients with *MAPK1*-related Rasopathy (MRR) exhibited craniofacial dysmorphism, intellectual disability, developmental delay, behavioral problems, small stature, hypotonia, and EEG abnormalities. Identified *MAPK1* mutations were classified as gain-of-function in functional studies since they enhanced ERK2 phosphorylation and nuclear translocation following growth factor stimulation *in vitro*^23^. However, pathogenic *MAPK1* variants have never been modeled in laboratory mammals, which would facilitate mechanistic studies to elucidate the pathophysiology of MRR and determine if ERK gain-of-function genocopies features observed in Rasopathy model mice with upstream Ras activation. To address this need, we generated the first mouse model of MRR to identify the physiological, anatomical, and neurodevelopmental sequelae of ERK2 gain-of-function *in vivo* and explore the role of downstream MAPK proteins in Rasopathy phenotypes.

## RESULTS

### Development of a mouse model of MAPK1-related Rasopathy

To model MRR, we used CRISPR to generate *Mapk1* mutant mice with an alanine-to-valine substitution at ERK2 position 172 (Figure 1A), which is equivalent to the human *MAPK1* A174V variant (c.521C>T; p.A174V) near the ERK2 activation segment that changes conformation when the kinase is phosphorylated at threonine 185 and tyrosine 187^23,24^. *Mapk1*^A172V/+^ crosses produced viable offspring for all genotypes (Figure 1B), and selective ERK2 activation (pERK2 and the pERK2:total ERK2 ratio) was confirmed in postnatal day 7 (PD7) liver samples (Figure 1C and Supplementary Figure 1) without a change in phosphorylated ERK1 or total ERK1/2 levels (Supplementary Figure 1). Of 476 mice genotyped before weaning, there was a trend toward reduced prevalence of mice carrying two copies of the mutant allele, but it did not deviate from Mendelian ratios (32.05% *Mapk1*^+/+^, 49.42% ; Chi-square test; *X*^2^ (2, N = 476) = 0.78, *p* = 0.68). In some mice, the *Mapk1*^A172V^ variant was associated with failure-to-thrive (body mass < 4.0 g), resulting in gene-dose-dependent postnatal mortality by PD21 (Figure 1D). This was most severe in *Mapk1*^A172V/A172V^ male mice, 52.63% of which died before reaching adulthood (Figure 1D, *middle*).

**Figure 1.**
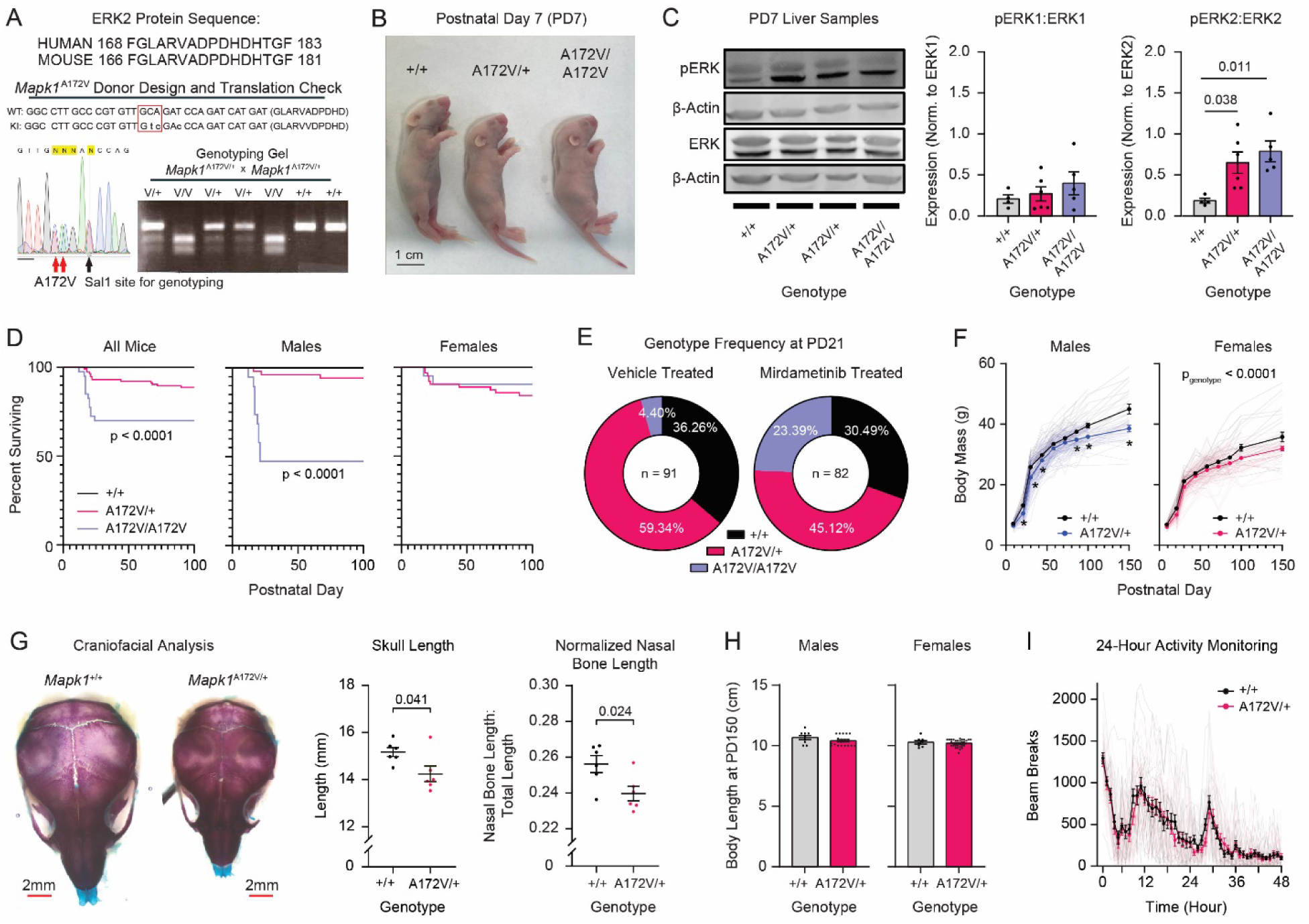
Development and characterization of a mouse model of *MAPK1*-related Rasopathy. **(A)** *Mapk1*^A172V^ transgenic mouse design showing homology between mouse and human sequences (*top*), sequencing confirmation of gene edit (bottom left), and a representative genotyping gel (*bottom right*). **(B)** *Mapk1*^+/+^, *Mapk1*^A172V/+^, and *Mapk1*^A172V/A172V^ pups at PD7 (scale bar: 1 cm). **(C)** Representative Western blot images of liver samples from *Mapk1* mutant and control mice (*left*). There were no differences in the pERK1:ERK1 ratio (*middle*) between genotypes (*n*_+/+_ = 4, *n*_A172V/+_ = 6, *n*_A172V/A172V_ = 5; one-way ANOVA: *F*2,12 = 0.84, *p* = 0.45). pERK2:ERK2 (*right*) was significantly increased (one-way ANOVA: *F*2,12 = 6.17, *p* = 0.014) in *Mapk1*^A172V/+^ (Bonferroni *post hoc* test; *p* = 0.038) and *Mapk1*^A172V/A172V^ mice (*p* = 0.011) relative to *Mapk1*^+/+^ controls. **(D)** *Mapk1*^A172V^ carriers exhibited significant early postnatal mortality (*left*; *n*_+/+_ = 61, *n*_A172V/+_ = 115, *n*_A172V/A172V_ = 40; log-rank test; *X*^2^(2, N = 216) = 23.66, *p* < 0.0001) that was largest in male *Mapk1*^A172V/A172V^ mice (*middle*; (*n*_+/+_ = 33, *n*_A172V/+_ = 52, *n*_A172V/A172V_ = 19; log-rank test; *X*^2^ (2, N = 104) = 42.40, *p* < 0.0001). No differences in mortality were observed in female *Mapk1* mutants (r*ight*; *n*_+/+_ = 28, *n*_A172V/+_ = 63, *n*_A172V/A172V_ = 21; log-rank test; *X*^2^ (2, N = 112) = 4.914, *p* = 0.086). **(E)** Low incidence of *Mapk1^A^*^172^*^V/A^*^172^*^V^*mice at PD21 was observed in vehicle-treated cages (*left*), deviating from Mendelian ratios (36.26% *Mapk1*+/+, 59.34% *Mapk1*A172V/+, and 4.40% *Mapk1*^A172V/A172V^ ; Chi-square: X (2, N = 91) = 15.79, p = 0.0004). Mirdametinib treatment (*right*) restored Mendelian ratios in *Mapk1* mutants (30.49% *Mapk1*+/+, 45.12% *Mapk1*^A172V/+^, and 24.39% *Mapk1*^A172V/A172V^ 0.57, p = 0.75). **(F)** Male (*left*) *Mapk1*^A172V/+^ mice have reduced body mass (mixed-effects analysis; *F*8,875 = 5.98, *p*day x genotype <0.0001; *F*_3.57,390.0_ = 1918, *p*day <0.0001; *F*1,147 = 31.45, *p*genotype < 0.0001), specifically at postnatal day 9, 21, 30, 44, 86, 100, and 150 (Bonferroni *post hoc* test; asterisk denotes *p* < 0.05). Female (*right*), *Mapk1*^A172V/+^ mice had lower mass than *Mapk1*^+/+^ littermates (mixed-effects analysis; *F*8,810 = 1.80, *p*genotype x day = 0.074; *F*1,138 = 16.20, *p*genotype <0.0001; *F*_3.42,345,9_ = 841.6, *p_day_* < 0.0001). **(G)** Stained skulls from *Mapk1*^A172V/+^ and *Mapk1*^+/+^ *A172V/+* mice mice at PD7 (*left*; scale bar: 2mm). *Mapk1^A^*^172^*^V/+^* had reduced skull length (*middle*; Mann-Whitney U-test; *U*(*n*_+/+_ = 6, *n*_A172V/+_ = 6) = 5, *p* = 0.041) and normalized nasal bone length (*right*; *n*_+/+_ = 6; *n*_A172V/+_ = 6; unpaired t-test; *t*10 = 2.65, *p* = 0.024). **(H)** No genotypic differences in body length were observed in male (*left*; *n*_+/+_ = 9; *n*_A172V/+_ = 19; t-test: *t*26 = 1.50, *p* = 0.15) and female (*right*; *n*_+/+_ = 8; *n*_A172V/+_ = 30; t-test: *t*36 = 0.63, *p* = 0.53) at PD150. **(I)** No differences in home cage locomotor activity was observed between genotypes (two-way repeated measures ANOVA; *F*_47,2115_ = 0.74, *p*_time x genotype_ = 0.90; *F*_1,45_ = 0.15, *p*_genotype_ = 0.52), but the test did reveal a significant main effect of time (*n*_+/+_ = 24; *n*_A172V/+_ = 23; two-way repeated measures ANOVA; *F*_12.87,579.2_ = 45.52, *p*_time_ < 0.0001) over the 48-hour period. Data is presented as mean ± standard error of the mean (SEM).

To investigate the cause of death, we collected blood and hearts at PD14. We did not observe cytopenia or leukocytosis (Supplementary Figure 2), which would be associated with a hematological or infectious cause of death, respectively. However, the relative distribution of leukocytes skewed towards the myeloid lineage in *Mapk1* mutants (Supplementary Figure 2), consistent with the known role of ERK in promoting myeloid cell differentiation^25^. Postnatal hearts in *Mapk1*^A172V/+^ and *Mapk1*^A172V/A172V^ mice displayed no lethal abnormalities (e.g. valvular, septal, large vessel, or myocardial defects; Supplementary Figure 3), which is consistent with MRR patients that experienced mild or no congenital heart defects^23^. Because MEK (the MAP kinase immediately upstream of ERK) inhibition reduced growth-factor stimulated ERK phosphorylation in MRR patient cells^23^, we hypothesized that the FDA-approved MEK inhibitor mirdametinib (PD0325901) would improve the survival of *Mapk1*^A172V/A172V^ mutants. Using the ‘MEK-in-milk’ protocol^26–28^, we administered mirdametinib to pups via the mother’s milk beginning 12 hours after birth (PD0.5) until PD21. In vehicle-treated cages, a high rate of mortality was observed in *Mapk1*^A172V/A172V^ pups, and the genotype frequency deviated from Mendelian ratios at PD21 (Figure 1E, *left*). However, no deaths occurred in mirdametinib-treated cages, restoring Mendelian ratios (Figure 1E, *right*) and confirming that downstream MAPK activation is pathological in mice modeling MRR.

Given that MRR patients carry a single pathogenic allele, we focused phenotyping efforts on *Mapk1*^A172V/+^ mice. *Mapk1*^A172V/+^ mice that survived past PD21 developed normally but had reduced body mass across their lifespan (Figure 1F), mirroring postnatal growth reduction in MRR patients^23^. In young mice, this was attributed to a reduction in body size (Figure 1B). For example, juvenile mutants had lower heart weight than controls that significantly correlated with body size (Supplementary Figure 3E). *Mapk1*^A172V/+^ mice also had smaller skulls (Figure 1G) without a change in aspect ratio (Supplementary Figure 4). Subsequent analysis of facial bone dimensions revealed a reduction in the nasal bone length in *Mapk1*^A172V/+^ mice that persisted after normalization to account for differences in skull size (Figure 1G, *right*; Supplementary Figure 4), recapitulating facial dysmorphism in NS mouse models^19,29–32^. In adult *Mapk1*^A172V/+^ mice, lower body mass was due to leaner body type rather than decreased body length (Figure 1H). This phenotype was not associated with differences in home cage activity level (Figure 1I) or steady state metabolism (Table 1), which has been observed in other Rasopathy models^32–34^. Thus, ERK2 activation is sufficient to recapitulate growth and craniofacial abnormalities observed in MRR patients and Rasopathy mouse models.

**Table 1.** Liver and plasma metabolites in *Mapk1* mutant and control mice. Metabolomic analysis of adult *Mapk1*^A172V/+^ and *Mapk1*^+/+^ mouse liver and plasma samples (n_+/+_ = 7, n_A172V/+_ = 6) showing steady state metabolite levels. Genotypic differences were examined using unpaired t-tests (p-values < 0.05 shown in bold). Data is presented as mean ± SEM.

### Neurodevelopmental effects of enhanced ERK2 signaling in Mapk1^A172V/+^ mice

Rasopathy patients may experience neurocognitive dysfunction, as well as abnormalities in brain morphology, white matter, and functional connectivity^35–39^. Therefore, we used a multidisciplinary approach to examine brain structure and function in *Mapk1*^A172V/+^ mice. First, we evaluated the time course of ERK phosphorylation changes in the forebrain samples from mutant and control mice (Figure 2A). On embryonic day 16.5 (ED16.5), ERK2 activation was higher in *Mapk1*^A172V/+^ forebrains relative to controls (Figure 2A, *left*) without a change in phosphorylated ERK1 or total ERK levels (Supplementary Figure 5). Likewise, at ED18.5, cortical and hippocampal expression of the transcription factor *Etv5* (Figure 2B), a downstream marker for MAP kinase pathway activation in the developing brain^40^, was expanded in *Mapk1*^A172V/+^ mice relative to *Mapk1*^+/+^ littermates. Increased forebrain ERK2 phosphorylation in *Mapk1*^A172V/+^ mice persisted postnatally and was largest at PD15 (Figure 2A, *middle*) but least robust in adulthood (Figure 2A, *right*). Phosphoproteomic analysis of PD15 forebrains identified 185 phosphopeptides with an abundance that was more than 1.5-fold different in *Mapk1*^A172V/+^ samples (71 lower, 114 higher) and had a p-value of less than 0.05 (Figure 2C). When genes associated with these phosphopeptides were used to identify differentially regulated biological pathways^41^, the top Reactome terms related to Ras/MAPK signaling or known functions of this pathway (e.g. *cellular responses to stress*, *cellular senescence*, *synaptic transmission*) (Figure 2D). To confirm that these changes related to ERK activation, we performed post-translational modification signature enrichment analysis (PTM-SEA)^42^ and observed that the ERK1/2 PTM-SEA score was significantly increased in *Mapk1*^A172V/+^ forebrains relative to wildtype controls (Figure 2E). Therefore, ERK activity is enhanced in the brains of mice modeling MRR.

**Figure 2.**
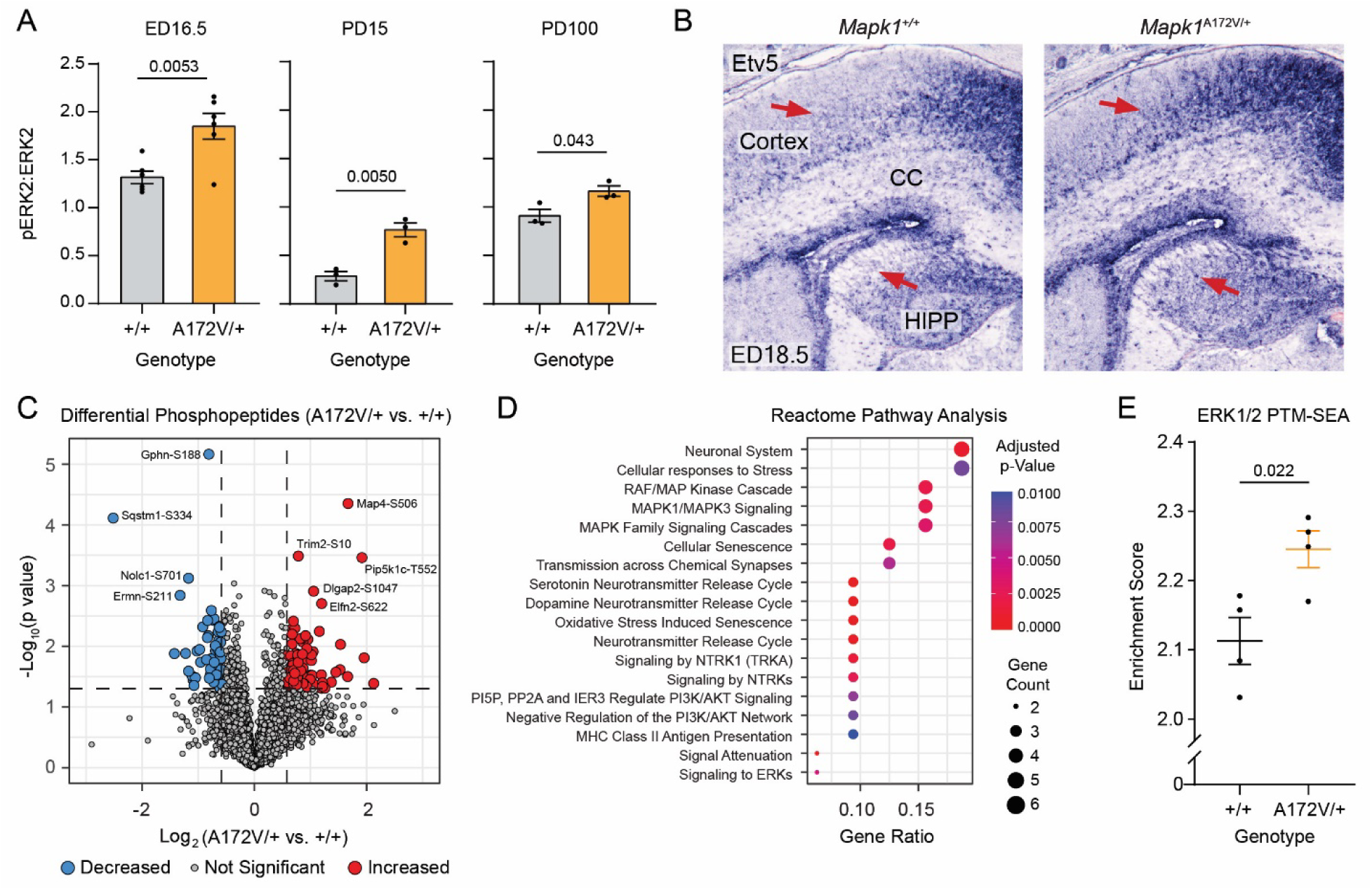
Proteomic characterization of ERK2 activity in the brains of *Mapk1*^A172V^ mice. **(A)** The pERK2:ERK2 ratio was increased in *Mapk1*^A172V/+^ mouse forebrain samples relative to controls at embryonic day 16.5 (ED16.5) (*left*; *n*_+/+_ = 6; *n*_A172V/+_ = 6; unpaired t-test; *t*10 = 3.55, *p* = 0.0053), PD15 (*middle*; *n*_+/+_ = 3; *n*_A172V/+_ = 3; unpaired t-test; *t*4 = 5.58, *p* = 0.0050), and PD100 (*right*; *n*_+/+_ = 3; *n*_A172V/+_ = 3; unpaired t-test; *t*4 = 2.93, *p* = 0.043). **(B)** Representative images showing *Etv5* expression in *Mapk1*^+/+^ and *Mapk1*^A172V/+^ mouse brains at ED18.5. Red arrows demonstrate expansion of *Etv5* expression in the cortex and hippocampus (HIPP) of *Mapk1*^A172V/+^ mice. CC = corpus collosum. **(C)** Volcano plot showing 5025 unique phosphopeptides that were detected in *Mapk1*^A172V/+^ and control forebrain samples (n = 4 per genotype) at PD15. Colored circles denote phosphopeptides > 1.5-fold increased (*red*) or decreased (*blue*) versus control with a p-value ≤ 0.05 (*horizontal dashed line*). **(D)** Reactome dot plot showing biological pathways significantly altered in *Mapk1*^A172V/+^ forebrain samples. **(E)** PTM-SEA analysis showed greater enrichment of ERK1/2 phosphorylation targets in *Mapk1*^A172V/+^ mice relative to *Mapk1*^+/+^ controls (*n*_+/+_ = 4; *n*_A172V/+_ = 4; unpaired t-test; *t* = 3.08, *p* = 0.022). Data is presented as mean ± SEM.

MAPK signaling is required for the generation of glial cells^43,44^, and dysregulated MAPK signaling alters their number and function in the brain^45–47^. For example, upstream MAPK activation by *Nf1* loss causes oligodendrocyte precursor cell (OPC) expansion, myelin decompaction, and reactive astrogliosis^48–52^. Therefore, we examined the effects of ERK2 gain-of-function on glial populations by staining PD14 brains for markers Pdgfrα, Nkx2.2, Myrf, and GFAP, which label OPCs, OPCs/early differentiating oligodendrocytes, mature oligodendrocytes, and reactive astrocytes, respectively. Like mouse models of NF1 and NS^49,51^, we observed an increase in the number of OPCs (Figure 3A-F) in the brains of *Mapk1*^A172V/+^ mice. While no difference in the number of Pdgfrα-positive cells (Figure 3A) was observed in the cerebral cortex (Figure 3B), OPC abundance was increased in the corpus collosum (Figure 3C) in mutant mice relative to controls. We also observed an increased number of Nkx2.2-positive OPCs/differentiating oligodendrocytes (Figure 3D) in both the cortex (Figure 3E) and corpus collosum (Figure 3F) in *Mapk1*^A172V/+^ mice without a change in the number of mature oligodendrocytes (Figure 3G-I). At 10 weeks of age, we examined mature white matter structure using transmission electron microscopy (Figure 3J). Like mice with oligodendrocyte-specific deletion of *Nf1*^50,53^, a gain-of-function *HRas* variant^54^, or mutant *Ptpn11*^51^, colossal white matter in *Mapk1*^A172V/+^ mice displayed evidence of myelin decompaction (Figure 3K). An expansion of GFAP expression, indicating reactive gliosis, was observed in subcortical structures, most prominently the diencephalon and hypothalamus, at this time point (Supplementary Figure 6). Thus, downstream MAPK activation via ERK2 gain-of-function causes glial abnormalities that genocopy those observed in mice with activating mutations upstream or at the level of Ras, such as *Nf1* loss or *Ptpn11*^Q79R^ expression.

**Figure 3.**
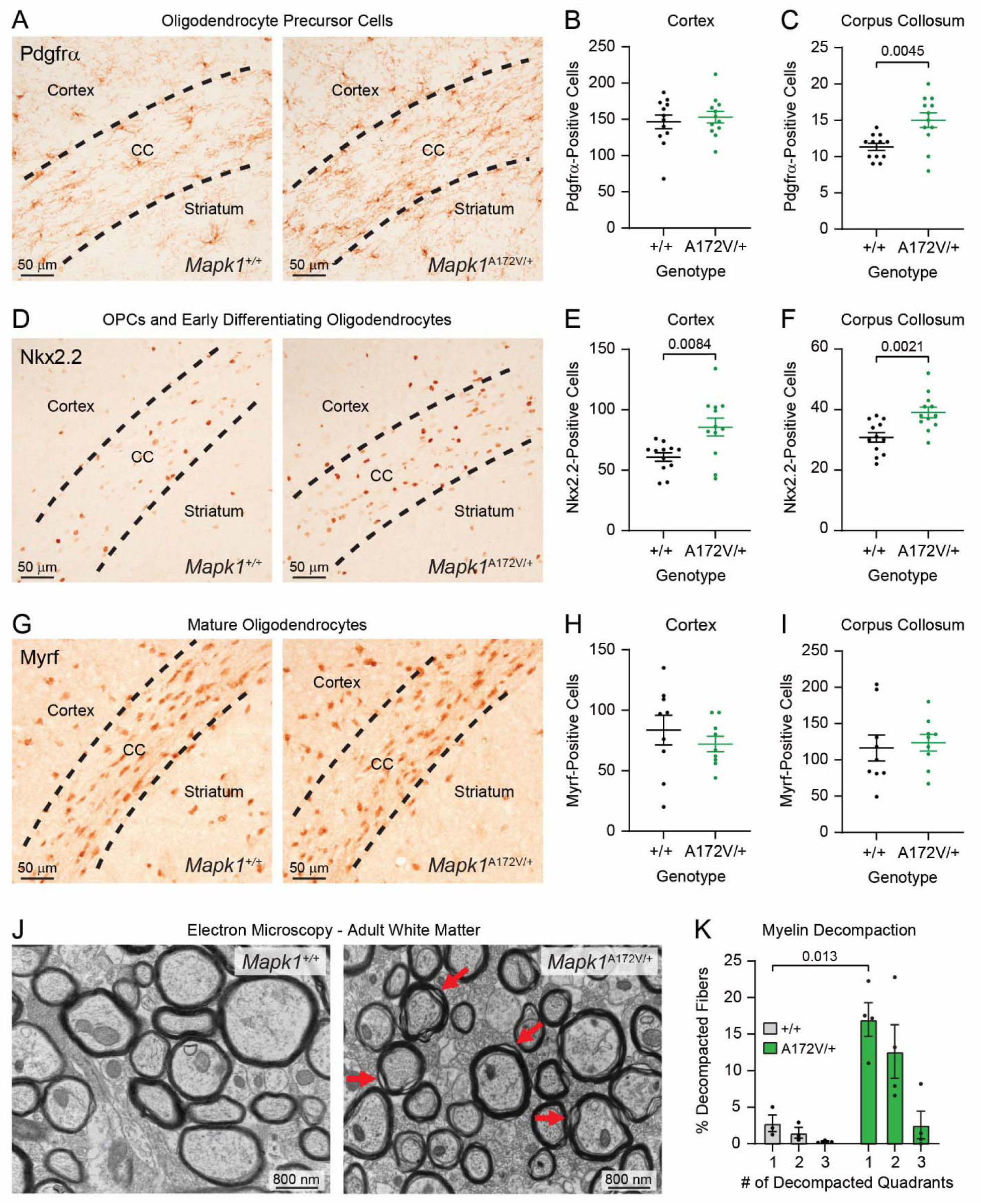
Examination of oligodendrocyte lineage defects in *Mapk1*^A172V/+^ mice. **(A)** Representative images showing Pdgfrα-stained oligodendrocyte precursor cells (OPCs) in PD14 *Mapk1*^+/+^ (left) and *Mapk1*^A172V/+^ mice (right). **(B-C)** There were no genotypic differences in the number of OPCs in the cerebral cortex (**B**; *n*_+/+_ = 12, *n*_A172V/+_ = 12; unpaired t-test; *t*_22_ = 0.51, *p* = 0.61), but more OPCs were observed in the corpus collosum (CC) in *Mapk1*^A172V/+^ brains relative to controls (**C**; *n*_+/+_ = 12, *n*_A172V/+_ = 12; unpaired t-test with Welch’s correction; *t*_15.85_ = 3.30, *p* = 0.0045). **(D)** Representative images of Nkx2.2-stained OPCs and differentiating early oligodendrocytes in *Mapk1*^+/+^ (*left*) and *Mapk1*^A172V/+^ mice (*right*). **(E-F)** Nkx2.2-positive cells were increased in the cortex (**E**; *n*_+/+_ = 12, *n*_A172V/+_ = 12; unpaired t-test with Welch’s correction; *t*_15.69_ = 3.012, *p* = 0.0084) and CC (**F**; *n*_+/+_ = 12, *n*_A172V/+_ = 12; unpaired t-test; *t*_22_ = 3.49 *p* = 0.0021) in *Mapk1*^A172V/+^ mice. **(G)** Representative images of Myrf-stained mature oligodendrocytes in *Mapk1*^+/+^ (*left*) and *Mapk1*^A172V/+^ mice (*right*). **(H-I)** There were no differences in Myrf-positive cells in the cortex (**H**; *n*_+/+_ = 9, *n*_A172V/+_ = 9; unpaired t-test; *t*_16_ = 0.83, *p* = 0.42) or CC (**I**; *n*_+/+_ = 9, *n*_A172V/+_ = 9; unpaired t-test; *t*_16_ = 0.35 *p* = 0.73). **(J)** Representative electron micrographs of corpus collosum samples from *Mapk1*^+/+^ (*left*) and *Mapk1*^A172V/+^ (*right*) mice at 10 weeks of age. Red arrows demonstrate areas of myelin decompaction. **(K)** There was a significant interaction between the number of decompacted quadrants and mouse genotype (*n*+/+ = 3, *n*A172V/+ = 4; two-way repeated measures ANOVA; *F*2,10 = 6.92, *p*_Decompacted Quadrants x_ Genotype = 0.013; *F*1.51,7.56 = 13.41, *p*Decompacted Quadrants = 0.0045; *F*1,5 = 11.69, *p*Genotype = 0.019), specifically in myelinated axons with 1 decompacted quadrant (Bonferroni *post hoc* test; *p* = 0.013). Scale bars in panels A, D, ang G are 50 μm. Scale bars in J are 800 nm. Data is presented as mean ± SEM.

### Mapk1^A172V/+^ mice recapitulate spatial memory deficits and sensory hypersensitivity observed in other Rasopathy mouse models

Patients with MRR may exhibit intellectual disability, ADHD, reduced stress tolerance, and aggressive behavior^23^. To determine the neurocognitive effects of ERK2 gain-of-function *in vivo*, we performed a battery of behavioral tests that assessed different aspects of neurological function in *Mapk1*^A172V/+^ and *Mapk1*^+/+^ littermates. While *Mapk1* mutant mice performed similarly to wildtypes in tests of motor function (e.g. open field locomotor activity, home cage activity monitoring, sucrose splash test; Supplementary Figure 7 A-B, Figure 1I), anxiety-like behavior (marble burying assay, dark/light box; Supplementary Figure 7C-D), short term memory (novel object recognition; Supplementary Figure 7E), and associative learning (contextual fear conditioning; Supplementary Figure 7F), unique phenotypes were identified. First, we observed that *Mapk1*^A172V/+^ mice – like other Rasopathy models^55–60^ – were slower to learn the location of a submerged platform using environmental cues in the Morris water maze, which assays spatial memory^61^. Specifically, they had longer latencies to the platform during the first two days of testing (the ‘training’ day when the platform was visible and acquisition day 1 when the platform was submerged and distal cues were present) (Figure 4A-B). With additional training, *Mapk1* mutants were able to achieve proficiency in the task, exhibiting no differences in platform latency during days 2-5 of the acquisition phase (Figure 4B) or during the probe trial (Supplementary Figure 7G), in which the platform is removed and the ability to remember its previous location is assessed. No differences in reversal learning were observed (Supplementary Figure 7H). Thus, enhanced ERK2 activity in the brain alters some but not all aspects of memory function and is sufficient to recapitulate Morris water maze deficits observed in mouse models of NF1 and NS.

**Figure 4.**
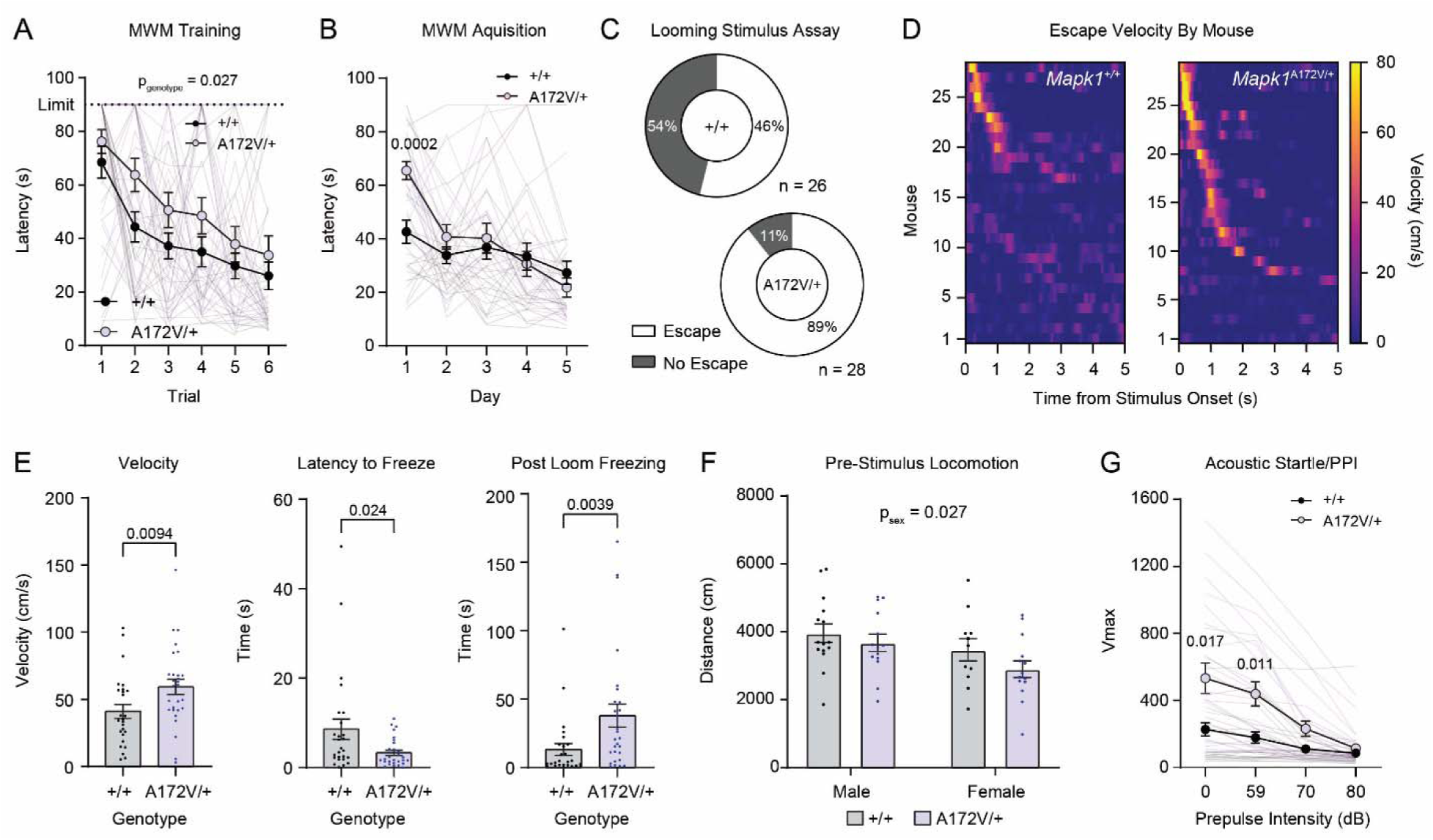
Behavioral characterization of *Mapk1*^A172V/+^ mice. **(A)** *Mapk1*^A172V/+^ mice had a longer latency to find the platform during Morris water maze (MWM) training (*n*+/+ = 25, *n*A172V/+ = 22; two-way repeated measures ANOVA; *F*5,225 = 0.38, *p*trial x genotype = 0.86; *F*1,45 = 5.25, *p*genotype = 0.027; *F*4.47,201.4 = 19.48, *p*trial < 0.0001). Four *Mapk1*^A172V/+^ mice failed to locate the platform and were excluded from further testing. **(B)** During the MWM acquisition phase, there was a significant day x genotype interaction (*n*+/+ = 25, *n*A172V/+ = 18; *F*4,164 = 5.44, *p* = 0.0004), and *Mapk1*^A172V/+^ mice had a longer latency to find the platform on the first acquisition day (Bonferroni *post hoc* test; *p* = 0.0002) compared to controls. **(C)** *Mapk1*^A172V/+^ mice had a higher probability of escape than *Mapk1*^+/+^ mice in the looming stimulus assay (*n*+/+ = 26, *n*A172V/+ = 28; Chi-square test; *X*^2^(2, N = 54) = 8.440, *p* = 0.0037). **(D)** Heat maps showing looming disc-evoked changes in velocity in individual *Mapk1*^+/+^ (*left*) and *Mapk1*^A172V/+^ mice (*right*). **(E)** *Mapk1*^A172V/+^ mice had a higher peak velocity (*left*; Mann-Whitney U-test, *U*(*n*+/+ = 26, *n*A172V/+ = 28) = 215, *p* = 0.0094), a shorter latency to freeze (*middle*; Mann-Whitney U-test, *U*(*n*+/+ = 26, *n*A172V/+ = 28) = 234.5, *p* = 0.024), and increased post loom freezing (*right*; Mann-Whitney U-test, *U*(*n*+/+ = 26, *n*A172V/+ = 28) = 199.5, *p* = 0.0039). **(F)** There were no differences in pre-stimulus locomotor activity between *Mapk1*^+/+^ mice and *Mapk1*^A172V/+^ mice during the looming stimulus assay; however, a significant decrease in distance travelled was observed in females independent of genotype (*n*+/+ = 26, *n*A172V/+ = 28; two-way ANOVA; *F*1,50 = 0.27, *p*sex x genotype = 0.60; *F*1,50 = 5.18, *p*sex = 0.027; *F*1,50 = 2.38, *p*genotype = 0.13). **(G)** There was a significant increase in the startle response of *Mapk1*^A172V/+^ mice relative to controls (*n* = 23, *n* = 23; two-way repeated measures ANOVA; *F*3,132 = 10.17, *p*stimulus x genotype = < 0.0001; *F*1.67,51.29 = 40.61, *p*stimulus < 0.0001; *F*1,44 = 8.45, *p*genotype = 0.0057). The startle response was significantly larger in response to the stimulus alone (Bonferroni *post hoc* test; *p* = 0.017) and in trials with the lowest intensity (59 dB) pre-pulse (Bonferroni *post hoc* test; *p* = 0.011). Data is presented as mean ± SEM.

Hypersensitivity to sensory stimuli is a common feature of autism and is prevalent in Rasopathy patients with co-morbid ASD^10,62^. Previously, we showed that mice modeling NF1 are more sensitive than wildtype littermates to arousing visual stimuli^63^, such as threatening looming discs that simulate aerial predator approach from above. In a looming stimulus assay, presentation of a train of expanding, dark, overhead discs on a light background promotes rapid flight to shelter followed by periods of freezing in rodents^64^. Like *Nf1*^+/-^ mice, *Mapk1*^A172V/+^ mice exhibited more robust behavioral responses to looming visual threats than littermate controls, including higher rates of escape (Figure 4C), increased post-stimulus velocity (Figure 4E, *left*), a shorter latency to freeze (Figure 4E, *middle*), and longer freezing duration (Figure 4E, *right*) in response to the train of looming discs without a difference in the distance travelled during the pre-stimulus habituation period (Figure 4F). To assay other sensory systems, we measured pre-pulse inhibition of the acoustic startle reflex in *Mapk1*^A172V/+^ and *Mapk1*^+/+^ mice. While sensory gating (i.e. pre-pulse inhibition) was intact in *Mapk1*^A172V/+^ mice, they exhibited more robust startle reflexes than littermate control mice when exposed to a loud (120 dB) acoustic stimulus (Figure 4G). This phenotype mirrors acoustic startle hypersensitivity previously observed in mice with biallelic *Nf1* loss in oligodendrocytes^50^. In addition to acoustic stimuli, *Mapk1* mutants had a shorter latency to groom in the sucrose splash test (Supplementary Figure 7B) without a change in the total grooming time, suggesting that they may be more sensitive to tactile stimuli. Therefore, ERK2 gain-of-function in *Mapk1*^A172V/+^ mice causes hypersensitivity to aversive sensory stimuli in mice, providing evidence that these phenotypes result from aberrant downstream MAPK signaling in mouse models of NF1 and other Rasopathy syndromes.

## DISCUSSION

In these studies, we generated the first genetically engineered mouse model of *MAPK1*-related Rasopathy (*Mapk1*^A172V/+^ mice), which recapitulates several clinical features observed in MRR patients: small stature, craniofacial dysmorphism, and cognitive impairment. Because the *Mapk1*^A172V^ pathogenic variant near the ERK2 activation domain increased kinase activity, we were able to address an important unanswered question in the Rasopathy field: to what extent do mice with gain-of-function mutations in downstream Ras effectors, like the terminal MAP kinase ERK, genocopy Rasopathy model mice with genetic perturbations acting upstream or at the level of Ras? We found that ERK2 gain-of-function is sufficient to produce a core set of central nervous system cellular, histological, and functional phenotypes observed in Rasopathy mouse models with upstream mutations: oligodendrocyte lineage defects, reactive astrocytosis, deficits in hippocampus-dependent spatial learning, and increased reactivity to arousing sensory stimuli, such as looming visual threats.

Consistent findings in mice modeling Rasopathy syndromes have been abnormalities in the generation and function of glial cell populations in the brain. Like mice modeling NF1^49,50,53^ and NS^51^, the brains of *Mapk1*^A172V/+^ mice had distinct oligodendrocyte lineage phenotypes with increased number of Nkx2.2- and Pdgfrα-positive OPCs and evidence of myelin decompaction in adulthood. We also found increased GFAP staining, an indicator for reactive gliosis which has also been observed in Rasopathy mouse models^48,51,65^. The Ras/MAPK pathway is required for gliogenesis, as deletion of MEK1/2 results in a cortex without astrocytes and oligodendrocytes, and hyperactivation of MEK1 increases astrocyte^43^ and OPC abundance^40^. Whereas *Ptpn11* loss-of-function reduces OPC generation, NS-associated *Ptpn11* mutants increase the number of OPCs and cause abnormal myelination^51^. To date, the ERK transcriptional targets driving aberrant oligodendrogenesis have not yet been identified. In Ras/MAPK mutant cancers, a conserved gene signature that includes *ETV4/5*, *SPRY2/4*, *DUSP4/6*, and *EPHA2/4* is a reliable biomarker of pathway activation and is associated with good clinical response to MEK inhibitor therapy^66^. Of these, the ETS transcription factor Etv5 is a strong candidate to play a mechanistic role in MRR-related deficits given its upregulation in *Mapk1*^A172V/+^ mice and involvement in the generation and maturation of neurons and glia in the hippocampus, cortex, and other brain regions^67–71^. Etv5 is also required for the maintenance of hippocampal neuron dendritic morphology and plasticity, as well as hippocampus-dependent learning^68^, in mice. Thus, future efforts will be required to explore the role of Etv5 and other ERK targets in OPC and cognitive deficits in MRR.

In addition to changes in OPC number during early postnatal development, *Mapk1*^A172V/+^ mice had evidence of myelin defects in mature white matter. Myelin decompaction has been observed in mice with conditional deletion of *Nf1*^50,53,72^ or expression of activated HRas^53,54^ or Shp2^51^ in oligodendrocytes. *Hras*^G12V^ or *Nf1*^−/-^ conditional mutants also have enhanced acoustic startle responses^50,53,72^, which raises the possibility that myelin decompaction in *Mapk1*^A172V/+^ mice causes sensory hypersensitivity phenotypes. In oligodendrocyte-targeted Rasopathy mouse models, myelination defects are MAPK-, nitric oxide-, and Notch-dependent^50,53,54^. Systemic treatment with the MEK inhibitor mirdametinib improved cell-autonomous white matter phenotypes in *Nf1* conditional knockout mice^50,53^, providing a roadmap for future mechanistic studies and/or preclinical therapeutic trials in MRR models. MEK inhibitors have an established safety profile, are FDA-approved for use in the treatment of NF1-associated tumors^73^, and prevented postnatal mortality in *Mapk1*^A172V/A172V^ mice. Dosing in *Mapk1* mutant mice will need to be carefully titrated, however, as mirdametinib treatment alone can perturb myelin structure in wildtype animals^54^, suggesting that the regulation of myelin integrity by MAPK signaling is tightly controlled.

*Mapk1*^A172V/+^ mice also exhibited deficits in hippocampus-dependent spatial learning in the Morris water maze. *Mapk1* mutants had longer trial times during initial training and increased latency to locate the platform on early acquisition trials. This result is consistent with phenotypes reported in other Rasopathy mouse models, including NS, Costello syndrome, and NF1^55–60,74,75^, as well as children with NF1 who participated in a virtual-reality Morris water maze task^76^. Previous studies in *Nf1*^+/-^ mice attributed learning deficits to impaired hippocampal long-term potentiation (LTP) caused by increased inhibitory GABA tone^55^ that occurs via ERK-mediated phosphorylation of synapsin I^56^. This presynaptic terminal protein controls the ready releasable pool of synaptic vesicles by regulating interactions with actin^77,78^. Additionally, Ras/MAPK signaling in the postsynaptic compartment plays an established role in hippocampal LTP^79–82^ by regulating AMPA receptor insertion^83,84^ and dendritic spine structural plasticity^85^. Thus, spatial learning deficits occurring in the context of *MAPK1* gain-of-function mutations are likely to have a complex neurophysiological etiology involving pre-synaptic regulation of GABAergic neurotransmission and/or changes in the post-synaptic compartment in glutamatergic pyramidal neurons. Future electrophysiological, molecular, and functional studies will be needed to test this hypothesis and clarify the role of different hippocampal cell types in spatial memory deficits in *Mapk1*^A172V/+^ mice.

Finally, we observed that *Mapk1*^A172V/+^ mice are hypersensitive to looming visual threats, a phenotype that we previously observed in *Nf1*^+/-^ mice^63^. In rodents, the ability of looming visual threats to rapidly evoke an innate defensive behavior is dependent on an evolutionarily conserved circuit involving the midbrain superior colliculus^86^, which is directly innervated by retinal ganglion cells in its superficial layers^87^. The saliency, or perceived importance, of a looming disc is encoded by the activity of glutamatergic neurons in the superior colliculus that, in turn, provide excitatory input to glutamatergic neurons in the dorsal PAG that compute escape decisions via a synaptic threshold mechanism^88^. The superior colliculus also innervates the midbrain ventral tegmental area, which can modulate looming stimulus-evoked escape via GABAergic projections to the central nucleus of the amygdala^89^ or dopaminergic projections to the nucleus accumbens medial shell^90^. Higher order brain regions like the retrosplenial^91^, visual^92^, and prefrontal^93^ cortices provide descending input to the superior colliculus that regulates different aspects of the escape behavior. At this time, it is unknown how supraphysiological MAPK signaling in subcortical sensory processing centers or the cortical sites that regulate them enhances threat reactivity in Rasopathy mice, which will need to be dissected in future studies.

## RESOURCE AVAILABILITY

### Lead Contact

Requests for more information should be directed to the lead contact, Dr. J. Elliott Robinson.

### Materials Availability

The *Mapk1^A^*^172^*^V^* mouse model is available upon request to the lead contact.

### Data and Code Availability

All source data used to produce the figures and tables, as well as the results of statistical testing procedures, are provided in the Source Data and Statistical Analysis file. Raw NMR and mass spectrometry data sets are available to download at: https://www.synapse.org/Synapse:syn71742374/. Code used for the looming stimulus assay is available at: https://github.com/jelliottrobinson/BonsaiLoomStim.

## ACKNOWLEDGEMENTS

This work was supported by NINDS project grant R01NS126108, SFARI Bridge to Independence Award 663007, CCHMC startup funds to JER; a CCHMC Center for Pediatric Genomics Pilot Award to RRW and CEP; NINDS training grant T32NS007453 to KEG; and shared instrumentation grant S10OD026717 to KDG.

## AUTHOR CONTRIBUTIONS

K.E.G.: data curation, formal analysis, investigation, project administration, visualization, validation, and original writing and editing of the manuscript. Z.R.C., A.V., E.M.C., K.S., C.A.B., and K.R.R.: data visualization, investigation, and formal analysis. A.A.F.: data curation, formal analysis, resources, software, and visualization. M.T.C., D.N., R.M.S., and A.W.: resources. T.A.R.: investigation and formal analysis. M.T.W.: formal analysis. C.V.V.: conceptualization. L.E.R., K.D.G., S.P.A., and K.N.W.: formal analysis and conceptualization. L.M.: conceptualization and resources. N.R.: conceptualization. C.E.P.: conceptualization, funding acquisition, and methodology. R.R.W.: conceptualization, funding acquisition, investigation, methodology, resources, and supervision. J.E.R.: conceptualization, funding acquisition, project administration, resources, supervision, visualization, writing of the original draft, and review and editing of the final manuscript.

## METHODS

### EXPERIMENTAL MODEL DETAILS

#### Mice

All experiments were conducted following the Institutional Animal Care and Use Committee (IACUC) of Cincinnati Children’s Hospital Medical Center (CCHMC) under protocols 2023-0044 and 2025-0007 and the National Institute of Health Guidelines for the Care and Use of Laboratory Animals. *Mapk1*^A172V^ mice were generated by the CCHMC Transgenic Animal and Genome Editing Core using CRISPR technology to introduce the A172V mutation plus a Sal1 site for genotyping. The correctly edited mutant allele founder mice (confirmed via Sanger sequencing) were bred to CD1 mice and maintained on this background. Heterozygous mutant mice (*Mapk1*^A172V/+^) were bred to produce wildtype, heterozygous, and homozygous mutant offspring. Mice were PCR genotyped with the following primers: Mapk1-5’-GTTTTCCTTGTTACTGATACTGCC and Mapk1-3’- ACTGAAGATGGTGACTCCTAAGC to amplify a 324 bp product which was subsequently digested with Sal1 restriction enzyme to produce 200bp and 124 bp products only in mutant mice. All mice were group housed by sex (up to four per cage) in standard shoebox cages on a 14/10 light/dark cycle with *ad libitum* access to food (LabDiet, Cat# 5010) and water. Mice were weighed at postnatal day (PD) 9, 21, 30, 44, 58, 72, 86, 100, and 150. Behavioral experiments were conducted after mice reached adulthood (after 12 weeks of age). Male and female mice were tested in all experiments. Marble burying, sucrose splash test, dark-light testing, novel object recognition, open field testing, and the looming stimulus assay were performed in the Robinson Lab at CCHMC. All other behavioral tests were performed at the Animal Behavior Core at CCHMC.

### METHOD DETAILS

#### Behavioral Assays

##### Marble burying

The protocol used was adapted from Angoa-Pérez et al^1^. Mice were introduced to a cage filled halfway with clean, leveled corncob bedding. In each cage, 15 opaque black marbles were arranged in an approximately equidistant 3 x 5 grid. After 30 minutes, the number of marbles buried was recorded. A marble was considered “buried’ if at least 2/3 of the marble was submerged. The mean number of marbles buried was calculated for each genotype.

##### Sucrose splash test

The protocol used was adapted from Frisbee et al^2^. Mice were sprayed with 10% sucrose solution on the upper neck between the ears and allowed to roam in a clean, empty cage for 5 minutes. Sessions were recorded using a Basler acA2040-120 um camera with an Edmunds Optics TECHSPEC 6mm C Series fixed focal length lens, and experimenters examined behavior every 5 seconds. Total grooming and latency to groom were calculated for each mouse.

##### Open field test

Recordings of each session were taken through a camera (Basler acA2040-120 um an Edmunds Optics TECHSPEC 6mm C Series fixed focal length lens). Mice were placed in the corner of a large plastic 55 cm x 45 cm storage container facing the wall and allowed to explore for 30 minutes. Ethovision^TM^ XT (Noldus Information Technology, Wageningen, the Netherlands) was used to determine the distance traveled and the time spent in the center of the arena.

##### Novel object recognition

The protocol used was adapted from Bevins and Besheer^3^. Specifically, in 55 cm x 45 cm arenas, two objects were placed equally spaced apart and fixed to the bottom of the container with Gorilla mounting putty. The objects were toy building blocks that were identical in size and color. Mice were allowed 10 minutes to explore the arena during the familiarization phase before being returned to their home cage. One hour later, the mice were returned to the same arena, but one of the objects was changed to a novel object, which was different in size and color from the original. The left-right location of the novel object was counterbalanced between mice. Again, mice were allowed to explore the objects for 10 minutes before being removed and placed back into their home cage. Videos for familiarization and novel object exploration were obtained with a Basler acA2040-120 um camera with an Edmunds Optics TECHSPEC 6mm C Series fixed focal length lens, and mouse position was tracked using Ethovision^TM^ XT software. Interaction with an object was determined when the nose of the mouse entered a region of interest that measured a circle with a 12 cm diameter around the object. The time of interaction was measured, and the discrimination ratio was calculated (time spent with novel object – time spent with familiar objects/total time interacting with both objects).

##### Dark-light box

The behavioral apparatus was a 37 cm x 20 cm arena that was one third covered by an opaque, infrared light (IR)-lucent shelter with an opening for mouse entry. The entire arena was exposed to bright light. Mice could move freely between dark and light sides. Behavior was recorded and tracked using Ethovision^TM^ XT. An IR light backlit panel was used under the arena to allow the software to detect the mouse inside of the shelter. Mice were placed in the lit side of the arena, facing toward the wall and away from the shelter. The latency to cross into the shelter, time spent in the shelter, time spent in the open arena, and crossings between the shelter and exposed arena were measured over a 10-minute period.

##### Looming

The looming stimulus assay was performed as described^4,5^. The clear acrylic arena measured 20.3 cm (w) x 61 cm (l) x 40.6 cm (h) and contained an (IR) light-lucent shelter at one side (the opening was the width of the arena) to serve as the escape location. A 9 cm petri dish was placed at the opposite end of the apparatus to encourage exploration of the ‘threat zone’ in the open arena. A 15.6-inch LCD monitor was placed above the threat zone to present a train of five looming stimuli (expanding black discs with a maximum diameter of 19.5 cm encompassing 27 degrees of visual angle) when the mouse entered the threat zone. The entire arena and monitor were placed inside a light and sound attenuating enclosure that could be closed during testing to minimize environmental stimuli during performance of the assay. An IR backlight (880 nm back-lit collimated backlight, Advanced Illumination) was used to allow mouse tracking in the light-attenuated chamber. To avoid overheating from the backlight, the arena was elevated on 4 small clear pedestals that were inaccessible to the mice, allowing airflow and space between the light and the floor of the arena. Mouse behavior was recorded using a Pylon camera with an Edmunds Optics TECHSPEC 6mm C Series fixed focal length lens positioned above the arena to allow for a full field of view. Ethovision^TM^ XT was used for real-time tracking and was synchronized with Bonsai^6^ to trigger the looming stimulus in the LCD monitor when the mouse entered the threat zone after a 10-minute habituation period. After the train of looming discs was presented, mice were recorded for 3 minutes before being returned to their home cage. Mice were excluded from analysis if they never entered the threat zone and thus never triggered a looming stimulus after 20 minutes.

##### Morris water maze

The arena was a pool that measured 150 cm in diameter and 51 cm high that contained a platform that was submerged 1-1.5 cm below the water level. The water was kept at ∼21°C. Mice were tracked using Anymaze software (Stoelting Company, Wood Dale, IL). There were 3 phases of the experiment: training, acquisition, and reversal learning. The starting quadrant was rotated in acquisition and reversal phases so no mouse started in the same quadrant on consecutive trials. Distance, speed, latency, and path efficiency were recorded for each phase; however, because of the coat color of the mice only latency was obtained. Training occurred on day 1 of testing. During this phase, curtains were drawn to cover external cues in the room. A 10 cm diameter platform with an orange ball was mounted to the platform. There was a 90-second time limit per trial with an inter-trial interval (ITI) of at least 10 minutes. If a mouse failed to find the platform at least twice within 6 trials, they were retested 4-6 hours later. Mice who found the platform moved on to the next phase of the experiment, while mice who failed to find the platform when retested were excluded (4 *Mapk1*^A172V/+^ mice). The acquisition phase occurred on days 2-7 with 5 days of learning and 1 memory (probe) trial on day 7. There were 4 trials each day with a 90-second limit per trial and an ITI of at least 10 minutes. During this phase, curtains were open to expose distal cues on the walls that mice could use to navigate the pool. The 10 cm diameter camouflaged platform was submerged 1-1.5 cm below the water surface. On the final day, during the probe trial, the platform was removed from the water bath to measure spatial memory for the platform on previous days. The final phase was reversal testing, which measures cognitive flexibility and was performed over 6 days of testing. The phase was identical to the acquisition phase, but a 7 cm diameter platform was placed into the opposite quadrant during the 5 days of learning. The platform was then removed during the probe trial on day 6.

##### Contextual fear conditioning

The experiment was conducted using the San Diego Instruments Freeze Monitor system (San Diego Instruments, San Diego, CA). The system contained a conditioning chamber with a metal bar floor for foot shock delivery (1.0 mA) and a photobeam array to record the start and end of freezing episodes. The test took place over two days. The first day was the habituation phase, which consisted of 5 minutes of acclimation followed by 9 intervals of shock delivered during the last 2 seconds of the 30-second ITI. This lasted 10 minutes. The second day was to test the mouse’s contextual memory. The mice were placed in the same chamber as day 1, but no foot shocks were delivered. This phase was 6 minutes long. Freezing behavior was measured in both phases.

##### 24 Hour home cage monitoring

The test was performed after P60 for two consecutive 24-hour recording periods using the HomeCage Photobeam Activity System (San Diego Instruments). Mice were individually housed in a static cage within the apparatus that contained a Napa Nectar gel and food pellets. After testing, mice returned to their home cage with their cage mates. Locomotor activity was measured as beam breaks per hour over 48 hours of testing.

##### Pre-pulse inhibition (PPI)

The test was performed in an SR-LAB apparatus (San Diego Instruments). Mice were placed in an acrylic cylindrical holder attached to an acrylic base plate with a piezoelectric accelerometer transducer to detect movement. During the session a fan was on, creating background ambient noise. Mice were given a 5-minute acclimation period before testing began. Pulses were 20-ms mixed frequency white noise bursts at 120 dB intensity. Pre-pulses were 59, 70, or 80 dB and were presented in counterbalanced order in a 20-trial block Latin Square. Peak response amplitude (Vmax) was recorded.

#### MEK Inhibitor in Milk Treatment

MEK inhibitor (MEKi) PD0325901 (Selleck Chemicals, Cat# 1036) was administered to lactating dams as described^7^. In brief, MEKi was dissolved in 0.5% methylcellulose and 0.2% Tween-80, which was used as the vehicle control. Cages were randomly selected for MEKi or vehicle treatment before pups were born, and MEKi (5mg/kg, which delivers∼1-2% of the maternal dose to pups via the milk^8^) or vehicle was delivered to lactating dams via oral gavage once a day from PD0.5 until PD21. Mice were euthanized for genotyping at PD21.

#### Blood Collection and Analysis

Trunk blood was collected from PD14 old mice into MiniCollect tubes with K2EDTA (Greiner Bio-One, Monroe, NC) following rapid decapitation. Samples were analyzed using a Heksa Element HT5 hematology analyzer to obtain a complete blood count with 5-part differential (Antech Diagnostics, Vermont, VIC, Australia). Samples were analyzed using a one-way ANOVA to determine differences between all 3 genotypes (*Mapk1*^+/+^*, Mapk1*^A172V/+^, and *Mapk1*A172V/A172V mice).

#### Tissue Extraction and Analysis

##### Brain protein extraction

Brains were extracted from E16.5, P15, and P100 mice following rapid decapitation. The cerebellum and hindbrain were removed and the forebrain was collected. One half of the forebrain was homogenized. RIPA Lysis Buffer (Thermo Fisher Scientific) and Halt Protease and Phosphatase Inhibitor Cocktail (Thermo Fisher Scientific) were added prior to homogenization. Then the tissue was agitated for 1 hour at 4°C and centrifuged at 14,000 xg for 14 minutes at 4°C. The supernatant was aliquoted and stored at -80°C until use. Protein was quantified using protocol 5.2 Microplate Assay Protocol of Detergent Compatible (DC) Protein Assay Instruction Manual (BioRad) and used for Western blots.

##### Western blot

Cell lysates were heated for 5 minutes at 95°C with 6X Laemmli SDS sample buffer, reducing (Thermo Fisher Scientific). Samples were loaded onto a 10% Mini-PROTEAN TGX Gel (BioRad) and run at 150 V until separated. Protein was transferred to a polyvinylidene difluoride membrane (PVDF) (BioRad) for 45 minutes at 100V. The PVDF membrane was blocked in Intercept Blocking Buffer (LI-COR) for 1 hour at room temperature, followed by incubation in primary antibody (pERK, Cell Signaling Technology, #4370, 1:1000; ERK, Cell Signaling Technology, #4696, 1:1000 or Cell Signaling Technology, #4695, 1:1000; and β-actin, Cell Signaling Technology, #3700, 1:1000) diluted in Intercept Antibody Diluent (LI-COR) at 4°C overnight. The following day, the membrane was washed 3 times for 5 minutes each with TBS-T before being incubated with the secondary antibody (IRDye 680RD Goat anti-Mouse IgG, LI-COR, #926-68070, 1:10,000 or 1:20,000; IRDye 800CW Donkey anti-Rabbit, LI-COR, #926-32213, 1:10,000 or 1:20,000) that was diluted in Intercept Antibody Diluent (LI-COR) for 1 hour at room temperature. The blot was then washed in TBS-T 3 times for 10 minutes followed by one wash for 10 minutes in TBS. Signals were visualized on the Odyssey CLx (LI-COR) using ImageStudio Ver 5.2 (LI-COR). Data was analyzed through Fiji and GraphPad Prism 10.1.2. If applicable, the PVDF membrane was stripped in Restore Western Blot Stripping Buffer (Thermo Fisher Scientific), followed by 30 minutes at room temperature in Intercept Blocking Buffer (LI-COR) before being incubated in the new primary antibody overnight at 4°C and repeating the second day protocol.

##### Phosphoproteomics

Phosphoproteomic quantification via mass spectrometry was performed at the UC Proteomics Laboratory at the University of Cincinnati College of Medicine using forebrain samples from P15 mice (4 *Mapk1*^+/+^ mice and 4 *Mapk1*^A172V/+^ mice, 2 males and 2 female samples per genotype). The brain tissue was frozen using liquid nitrogen directly in Kimble Kontes homogenizer tubes (item# 749520-0000), broken up within the tube with a small spatula, and homogenized in urea lysis buffer, which contained: 9 M urea, 20 mM HEPES, 1 mM b-glycero-phosphate, 2.5 mM sodium pyrophosphate, and 1 mM activated sodium orthovanadate. The samples were adjusted to 1 mL total volume and probe sonicated at 15 W for 15 seconds for 3 cycles. Unsolubilized material was pelleted via centrifugation, and samples were placed in new tubes. 3 mg aliquots were removed from each sample, and the volume was adjusted up to 1 mL with undiluted urea lysis buffer. The samples were digested with modified trypsin (Worthington Biochemical Corp., Cat# LS003740) overnight at 37°C. The digested peptides were purified and concentrated by passing them over a Sep-Pak C18 cartridge (Waters Corp., Cat# WAT051910), dried in a SpeedVac, and resuspended in binding/equilibration buffer from the Thermo Fisher High-Select TiO_2_ Phosphopeptide Enrichment Kit (Cat# A39223), which was used for peptide enrichment according to the manufacturer instructions. Eluted samples were dried in a SpeedVac and resuspended in 0.1% formic acid. 10% of each enriched sample was analyzed by nanoscale liquid chromatography coupled with tandem mass spectrometry (Orbitrap Eclipse).

All downstream analysis of the mass spectrometry data was performed using R v.4.5.0. Phosphosites were filtered for a high FDR and a Site Localization Score (ptmRS module) greater than 0.75 and unambiguous. The *PhosR* package was used for filtering, imputation, and scaling. Phosphopeptides with greater than 25% missing values across all samples were filtered out. For remaining phosphopeptides, site and condition specific imputation was performed if at least 50% of a given phosphopeptide was present in the defined condition (*Mapk1*^+/+^ and *Mapk1*^A172V/+^ WT). Differentially abundant phosphopeptides were identified using a t-test (p <0.05) and a fold change cutoff of |1.5|. The following packages were used for subsequent downstream analysis: *ReactomePA* (Reactome Pathway analysis), *ggplot2* (bargraphs and volcano plot), *ggpubr* (stats on bargraphs), and *ssGSEA2* (post-translational modification-signature enrichment analysis or PTM-SEA). For the PTM-SEA analysis, the ptm.sig.db.all.uniprot.mouse.v2.0.0 database was used, with all other parameters set to their default states.

##### NMR Metabolomics

The NMR based metabolomics experiments were performed at Cincinnati Children’s Hospital Medical Center Translational Metabolomics Facility (RRID: SCR_022636). Liver tissue samples were extracted using modified Bligh and Dyer methods^9–11^ to obtain polar metabolites. The extraction solvent volumes were determined based on the average water content of mouse liver tissue (72.8%). Plasma samples were processed according to the published protocol^12^. Briefly, the plasma samples were filtered through pre-washed 3 kDa spin filters and the filtrate was mixed with NMR buffer up to the final volume of 220uL. NMR data collection and processing were performed with a Bruker Avance III HD 600 MHz spectrometer using Topspin 3.6 software (Bruker Analytik, Rheinstetten, Germany), as previously described^13^. Total of 49 liver and 40 plasma metabolites were assigned based on the chemical shifts on 1D 1H-NOESY, 2D-TOCSY and -HSQC NMR experiments with reference spectra found in databases, Human Metabolome Database (HMDB)^14^, and Chenomx® NMR Suite profiling software (Chenomx Inc. version 8.1). The metabolites were quantified by Chenomx software based on the internal standard, TMSP and were normalized to tissue weights or plasma volume prior to statistical analysis by R studio and MetaboAnalyst^15^. Log transformation and mean centering data scaling were applied prior to multivariate analysis in MetaboAnalyst.

#### Immunohistochemistry

##### Tissue Collection and Cryostat Sectioning

Mice were transcardially perfused with PBS followed by 4% paraformaldehyde (PFA). The brain was extracted into PFA at 4°C overnight. The following day, the brains were transferred to 20% sucrose for cryoprotection. Brains were mounted in Tissue-Tek O.C.T Compound (Thermo Fisher Scientific) and sectioned in the cryostat at -20°C into 12 µm serial sections and mounted on slides. Slides were stored at -80°C until use.

##### Immunohistochemistry

In Coplin jars, warmed citrate buffer was poured over slides for antigen retrieval for a maximum of 5 minutes. Slides were then bleached with 0.33% H_2_O_2_ for 10 minutes and washed three times for 5 minutes each with 1X KPBS. Slides were then incubated in primary antibodies overnight in a humid chamber (Nkx2.2, Abcam, #ab191077, 1:1000; GFAP, Agilent, #Z0334, 1:2000; PDGFRα, R&D Systems, #AF1062, 1:1000, and Myrf, ABclonal, #A16355, 1:1000). The following day, slides were washed in 1X KPBS for 5 minutes three times. A biotinylated secondary antibody was added (Biotin-SP (long spacer) AffiniPure® Donkey Anti-Goat IgG (H+L), Jackson ImmunoResearch, #705-065-147, 1:500; Biotin-SP (long spacer) AffiniPure® Donkey Anti-Rabbit IgG (H+L), Jackson Immuno, # 711-065-152, 1:500) for 2 hours at room temperature in a humid chamber. Slides are washed three times for 5 minutes each in 1X KPBS and incubated in Avidin-Biotin Complex (ABC) (Vector Laboratories) for 1 hour in a humid chamber at room temperature. Slides are washed again three times for 5 minutes in 1X KPBS. Diaminobenzidine (DAB) is then added to slides in a humid chamber for 5 minutes, then washed in 1X KPBS three times for 5 minutes. Slides are left to dry overnight before being coverslipped. To coverslip, slides are rehydrated for 5 minutes before dehydrated in a series of ETOH washes (70%, 95%, and 100%) for 5 minutes each. Slides are then put into two washes of xylene for 5 minutes each and cover slipped with DPX mountant media (Sigma-Aldrich). CellProfiler (ver 4.2.8) was used to quantify the images.

##### Heart Immunohistochemistry

Histological analysis was conducted on formalin-fixed tissues, paraffin-embedded, sectioned at 5-8 μm, and stained either with hematoxylin and eosin (H&E) or with Russell-Movat pentachrome stain^16^. Whole-heart H&E-stained sections were viewed on a Zeiss Discovery V8 stereomicroscope and imaged with an Axiocam 820 camera. Valve pentachrome-stained sections were viewed on a Zeiss Axio Imager Z1 with an Axiocam 305 camera. Images were viewed and measured using Zeiss ZEN Blue software. The myocardial thickness for each sample was measured at three positions for each ventricle: the dorsal, lateral, and ventral surface. Equal number of sections were selected at random for each set of comparisons.

##### Skull Skeletal Preps

Heads from PD7 *Mapk1*^+/+^ and *Mapk1*^A172V/+^ pups were deskinned and fixed in ethanol for 2 days. Skulls were then washed in Alcian blue dye (Sigma Aldrich) overnight at room temperature to stain for cartilage. Skulls were rinsed with ethanol for two days and cleared with 1% KOH for 6 hours until clear. After clearing, skulls were counterstained with Alizarian red stain (Sigma Aldrich) for 24 hours to visualize bone. Finally, skulls were cleared with fresh 20% glycerol/1% KOH solution daily until the clearing was complete and stored in a 50% glycerol/50% ethanol solution. Skulls were imaged using a Zeiss Discovery V8 stereomicroscope with an Axiocam 820 color camera and ZEN Blue software.

### QUANTIFICATION AND STATISTICAL ANALYSIS

All statistical tests were conducted using R or GraphPad Prism version 10.5 (GraphPad). Sex was considered as a variable in a two-way or three-way ANOVA when appropriate. If no sex differences were found, the statistical testing was performed with the sexes combined. Parametric tests were conducted unless the data was found to be nonparametric using the D’Agostino-Pearson test for normality. For data that had small sample sizes, we used the Shapiro-Wilk test of normality instead. A Welch’s correction was used for t-tests if sample variances were significantly different. For body mass data, a mixed-effects model with the Geisser-Greenhouse correction was used instead of a repeated-measures ANOVA to account for missing values. The Bonferroni correction was used for *post hoc* testing after an ANOVA if any variable had three or more levels. A Dunn’s *post hoc* test was used for Kruskal-Wallis tests, and the Fisher’s Least Significant Difference (LSD) test was used for all other *post hoc* tests. Inheritance data was calculated using a Chi-square test of independence. Survival data was calculated using the log-rank test and plotted using a Kaplan-Meier plot. For correlations, we used Pearson’s correlation coefficient to determine the strength of a relationship between two variables, and linear regressions were run to generate regression lines on the graphs. All source data used to produce the figures and the results of statistical testing procedures are provided in the Source Data and Statistical Analysis file.

