## Supplementary Figures and Captions for "Genetic activation of ERK2 recapitulates core neurodevelopmental features of Rasopathy syndromes in mice"

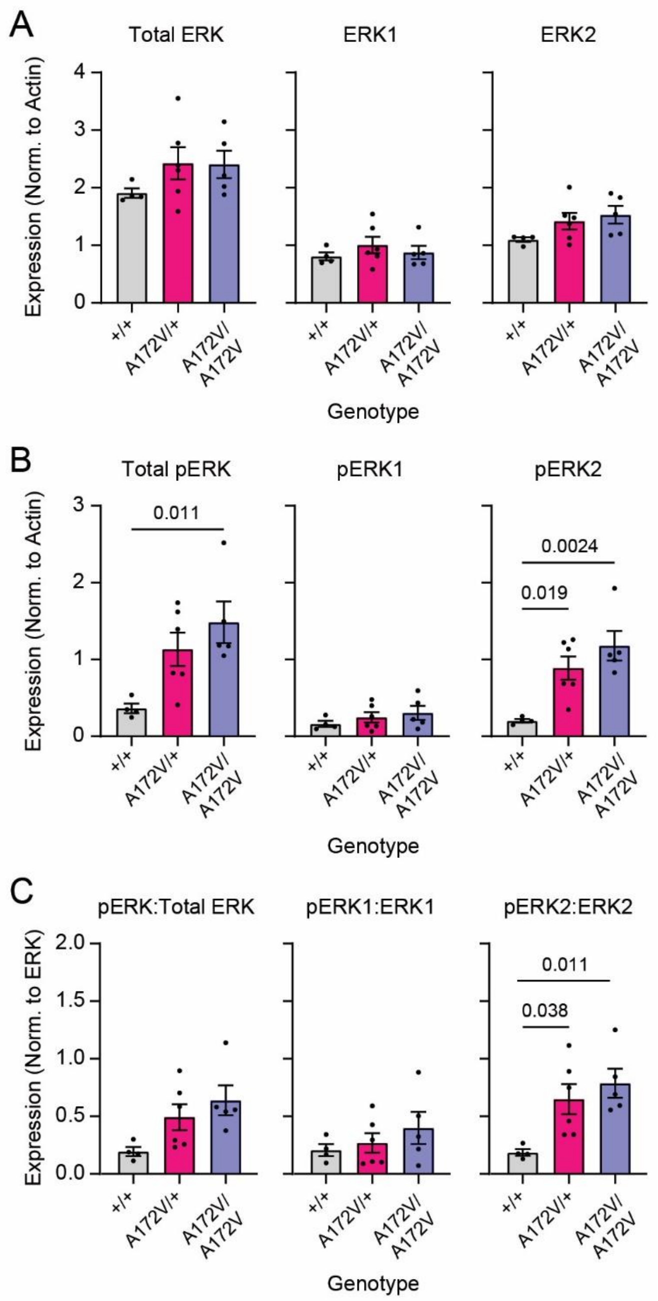


**Supplementary Figure 1. Analysis of ERK phosphorylation in PD7 livers in *Mapk1* mutant and control mice. (A)** There were no significant genotypic differences in total ERK (*left*; $n_{+/+}$ = 4; $n_{A172V/+}$ = 6; $n_{A172V/A172V}$ = 5; one-way ANOVA; $F_{2,12}$ = 1.28, $p$ = 0.31), ERK1 (*middle*; $n_{+/+}$ = 4; $n_{A172V/+}$ = 6; $n_{A172V/A172V}$ = 5; one-way ANOVA; $F_{2,12}$ = 0.65, $p$ = 0.54), or ERK2 (right; $n_{+/+}$ = 4; $n_{A172V/+}$ = 6; $n_{A172V/A172V}$ = 5; one-way ANOVA; $F_{2,12}$ = 2.37, $p$ = 0.14) in PD7 liver samples, as measured by

Western blot. **(B)** There was a significant increase in total pERK expression in PD7 *Mapk1* mutant livers (*left*; $n_{+/+}$ = 4; $n_{A172V/+}$ = 6; $n_{A172V/A172V}$ = 5; one-way ANOVA; $F_{2,12}$ = 5.95, $p$ = 0.016). Bonferroni *post hoc* analysis showed that pERK was significantly lower in *Mapk1*$^{+/+}$ samples relative to *Mapk1*$^{A172V/A172}$ samples ($p$ = 0.011); this difference did not reach statistical significance in *Mapk1*$^{A172V/+}$ livers relative to control samples ($p$ = 0.064). There were no genotypic differences in pERK1 (*middle*; $n_{+/+}$ = 4; $n_{A172V/+}$ = 6; $n_{A172V/A172V}$ = 5; one-way ANOVA; $F_{2,12}$ = 0.85, $p$ = 0.45). pERK2 levels were higher in *Mapk1* mutant livers relative to control samples (*right*; $n_{+/+}$ = 4; $n_{A172V/+}$ = 6; $n_{A172V/A172V}$ = 5; one-way ANOVA; $F_{2,12}$ = 9.18, $p$ = 0.0038). Bonferroni *post hoc* tests showed a significant increase in heterozygous mutants ($p$ = 0.019) and homozygous mutants ($p$ = 0.0024). **(C)** There were no genotypic differences in the pERK:total ERK ratio (*left*; $n_{+/+}$ = 4; $n_{A172V/+}$ = 6; $n_{A172V/A172V}$ = 5; one-way ANOVA; $F_{2,12}$ = 3.65, $p$ = 0.058) or pERK1:ERK1 ratio (*middle*; $n_{+/+}$ = 4; $n_{A172V/+}$ = 6; $n_{A172V/A172V}$ = 5; one-way ANOVA; $F_{2,12}$ = 0.84, $p$ = 0.46). The pERK2:ERK2 ratio was significantly increased in *Mapk1* mutants relative to wildtype controls (*right*; $n_{+/+}$ = 4; $n_{A172V/+}$ = 6; $n_{A172V/A172V}$ = 5; one-way ANOVA; $F_{2,12}$ = 6.17, $p$ = 0.014). pERK2:ERK2 was higher in *Mapk1*$^{A172V/+}$ (Bonferroni *post hoc* test; $p$ = 0.038) and *Mapk1*$^{A172V/A172V}$ livers (Bonferroni *post hoc* test; $p$ = 0.011) relative to *Mapk1*$^{+/+}$ samples. Data is presented as mean ± SEM.

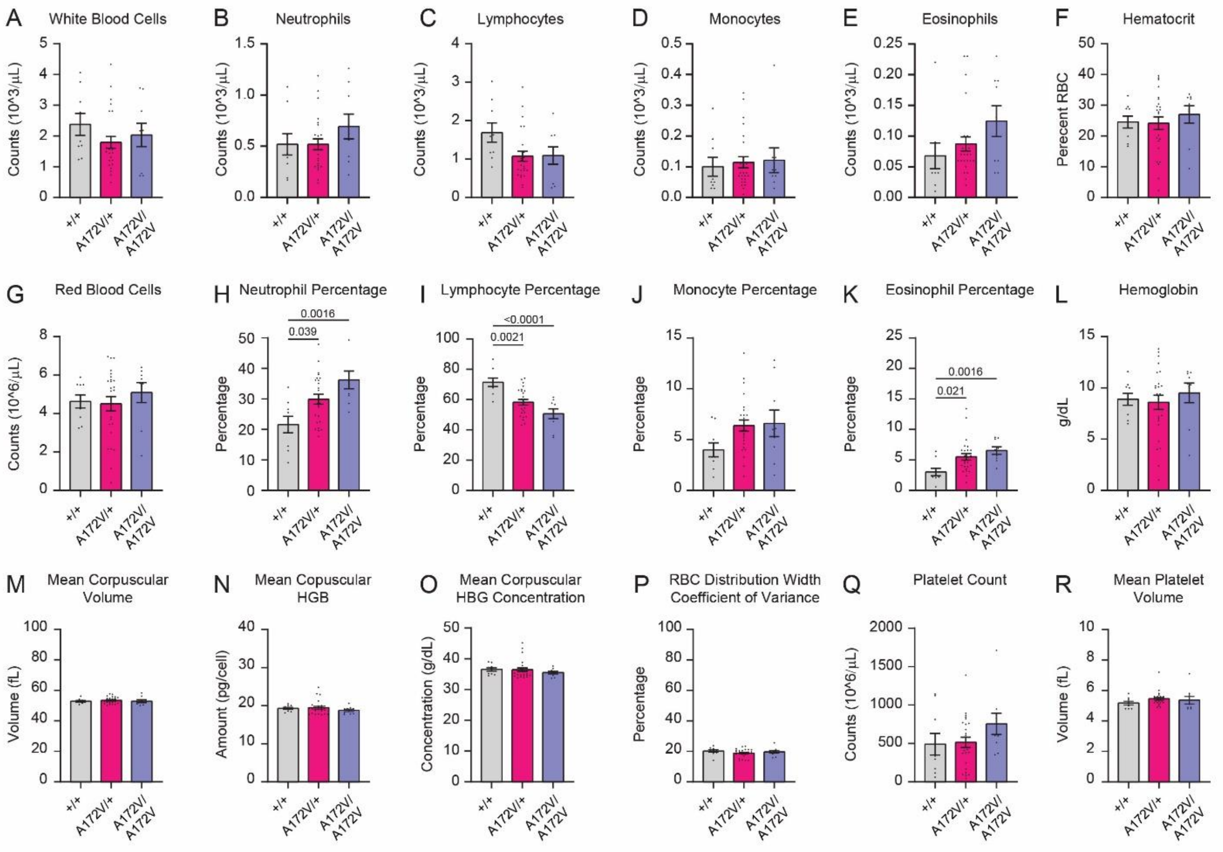


**Supplementary Figure 2. Hematological testing in *Mapk1* mutant and control mice. (A)** At two weeks of age, there were no significant differences in white blood cell (WBC) counts between in *Mapk1* mutant and control mice ($n_{+/+}$ = 9; $n_{A172V/+}$ = 25; $n_{A172V/A172V}$ = 9; one-way ANOVA; $F_{2,40}$ = 0.45, $p$ = 0.35). **(B-E)** There were no genotypic differences in absolute neutrophil ($n_{+/+}$ = 9; $n_{A172V/+}$ = 25; $n_{A172V/A172V}$ = 9; one-way ANOVA; $F_{2,40}$ = 0.61, $p$ = 0.31), lymphocyte ($n_{+/+}$ = 9; $n_{A172V/+}$ = 25; $n_{A172V/A172V}$ = 9; one-way ANOVA; $F_{2,40}$ = 0.27, $p$ = 0.069), monocyte ($n_{+/+}$ = 9; $n_{A172V/+}$ = 25; $n_{A172V/A172V}$ = 9; Kruskal-Wallis test; $H_3$ = 1.26, $p$ = 0.53), or eosinophil counts ($n_{+/+}$ = 9; $n_{A172V/+}$ = 25; $n_{A172V/A172V}$ = 9; Kruskal-Wallis test; $H_3$ = 2.82, $p$ = 0.24). **(F-G)** There were no genotypic differences in hematocrit ($n_{+/+}$ = 9; $n_{A172V/+}$ = 25; $n_{A172V/A172V}$ = 9; one-way ANOVA; $F_{2,40}$ = 0.33, $p$ = 0.72) or red blood cell (RBC) count ($n_{+/+}$ = 9; $n_{A172V/+}$ = 25; $n_{A172V/A172V}$ = 9; one-way ANOVA; $F_{2,40}$ = 0.41, $p$ = 0.67). **(H)** There was a significant increase in the percentage of neutrophils in *Mapk1* mutants ($n_{+/+}$ = 9; $n_{A172V/+}$ = 25; $n_{A172V/A172V}$ = 9; one-way ANOVA; $F_{2,40}$ = 7.21, $p$ = 0.0021).

Bonferroni *post hoc* analysis revealed a significant increase in both *Mapk1*$^{A172V/+}$ ($p$ = 0.039) and *Mapk1*$^{A172V/A172}$ ($p$ = 0.0016) neutrophil percentage compared to wild-type littermates. **(I)** There was a significant decrease in the percentage of lymphocytes ($n_{+/+}$ = 9; $n_{A172V/+}$ = 25; $n_{A172V/A172V}$ = 9; one-way ANOVA; $F_{2,40}$ = 11.87, $p$ < 0.0001) in *Mapk1*$^{A172V/+}$ (Bonferroni *post hoc* test; $p$ = 0.0021) and *Mapk1*$^{A172V/A172V}$ (Bonferroni *post hoc* test; $p$ < 0.0001) mice. **(J)** There was no genotypic difference in monocyte percentage ($n_{+/+}$ = 9; $n_{A172V/+}$ = 25; $n_{A172V/A172V}$ = 9; one-way ANOVA; $F_{2,40}$ = 2.52, $p$ = 0.093). **(K)** There was a significant increase in the percentage of eosinophils ($n_{+/+}$ = 9; $n_{A172V/+}$ = 25; $n_{A172V/A172V}$ = 9; Kruskal-Wallis test; $H_3$ = 12.57, $p$ = 0.0019) in both *Mapk1*$^{A172V/+}$ mice (Dunn's *post hoc* test; $p$ = 0.021) and *Mapk1*$^{A172V/A172V}$ mice (Dunn's *post hoc* test; $p$ = 0.0016). **(L-O)** There were no genotypic differences in hemoglobin (HGB) concentration ($n_{+/+}$ = 9; $n_{A172V/+}$ = 25; $n_{A172V/A172V}$ = 9; one-way ANOVA; $F_{2,40}$ = 0.30, $p$ = 0.74), mean corpuscular volume ($n_{+/+}$ = 9; $n_{A172V/+}$ = 25; $n_{A172V/A172V}$ = 9; one-way ANOVA; $F_{2,40}$ = 0.38, $p$ = 0.68), mean corpuscular HGB ($n_{+/+}$ = 9; $n_{A172V/+}$ = 25; $n_{A172V/A172V}$ = 9; Kruskal-Wallis test; $H_3$ = 1.24, $p$ = 0.54) or corpuscular HBG concentration ($n_{+/+}$ = 9; $n_{A172V/+}$ = 25; $n_{A172V/A172V}$ = 9; Kruskal-Wallis test; $H_3$ = 1.58, $p$ = 0.45). **(P)** There were no genotypic differences in the coefficient of variation of RBC distribution width ($n_{+/+}$ = 9; $n_{A172V/+}$ = 25; $n_{A172V/A172V}$ = 9; one-way ANOVA; $F_{2,40}$ = 0.89, $p$ = 0.42). **(Q-R)** There were no genotypic differences in platelet count ($n_{+/+}$ = 9; $n_{A172V/+}$ = 25; $n_{A172V/A172V}$ = 9; one-way ANOVA; $F_{2,40}$ = 1.60, $p$ = 0.22) or average volume ($n_{+/+}$ = 9; $n_{A172V/+}$ = 25; $n_{A172V/A172V}$ = 9; Kruskal-Wallis test; $H_3$ = 3.82, $p$ = 0.15). Data is presented as mean ± SEM.

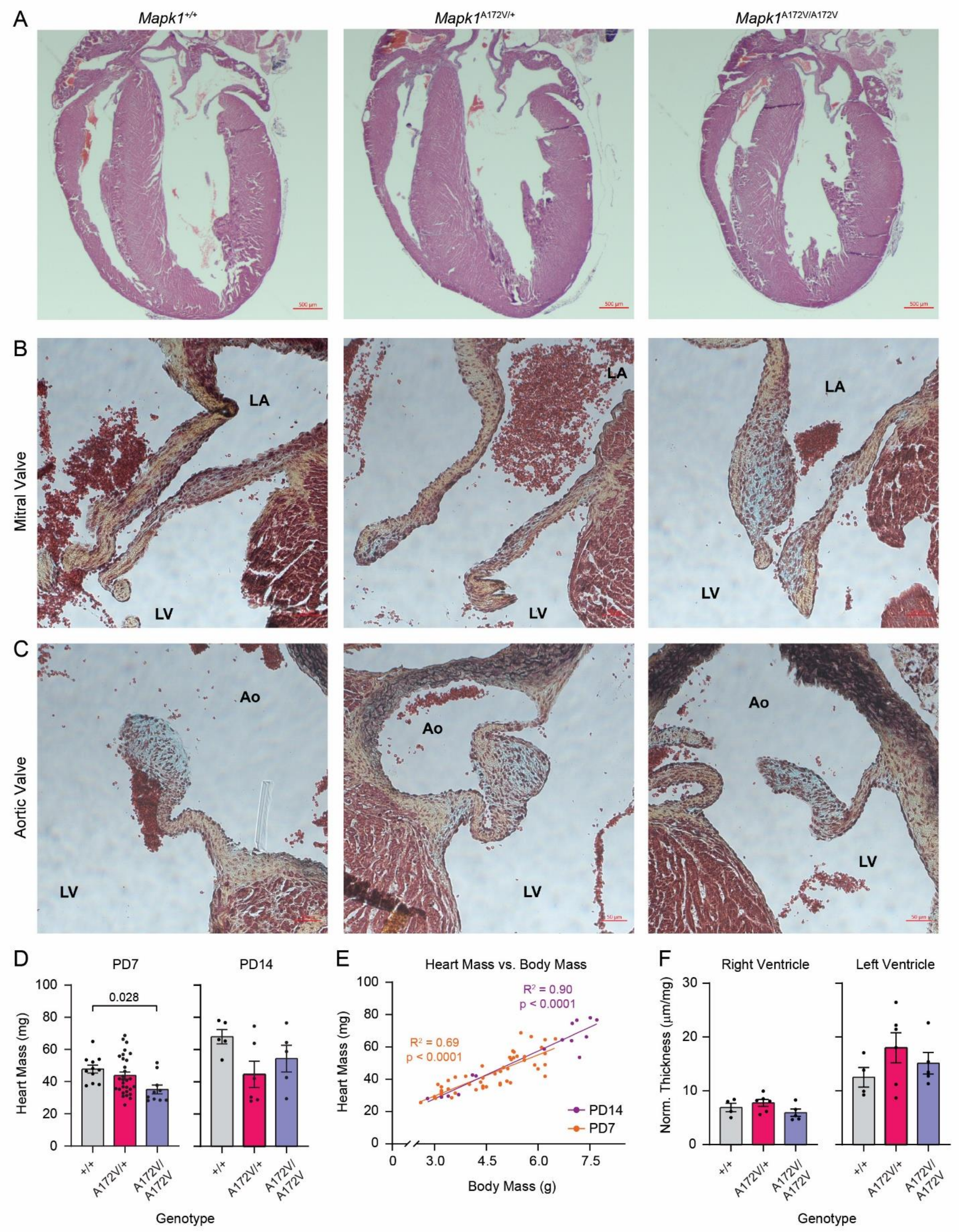


**Supplementary Figure 3. Myocardial and valvular analysis of *Mapk1* mutant and control**

**hearts. (A-C)** Representative cross-sectional images of the myocardium (**A**), mitral valve (**B**), and aortic valve **(C)** from 2-week-old *Mapk1*$^{+/+}$ (*left*), *Mapk1*$^{A172V/+}$ (*middle*), and *Mapk1*$^{A172V/A172V}$ mice (*right*) hearts. **(D)** There was a significant difference in the heart weight of *Mapk1* mice at PD7 (*left*; $n_{+/+}$ = 11, $n_{A172V/+}$ = 30, $n_{A172V/A172V}$ = 10; one-way ANOVA; $F_{2,48}$ = 3.89, $p$ = 0.027) with a significant reduction in *Mapk1*$^{A172V/A172V}$ heart mass compared to wild-type littermates (Bonferroni *post hoc* test; $p$ = 0.028). Genotypic differences in heart mass at PD14 did not reach statistical significance (*right*; $n_{+/+}$ = 5, $n_{A172V/+}$ = 6, $n_{A172V/A172V}$ = 5; one-way ANOVA; $F_{2,13}$ = 2.57, $p$ = 0.11). **(E)** There was a significant correlation between heart mass and body mass at PD7 ($R^2(49)$ = 0.69, $p$ < 0.0001) and PD14 ($R^2(13)$ = 0.90, $p$ < 0.0001). **(F)** There was no significant change in normalized right ventricular (one-way ANOVA; $F_{2,12}$ = 1.82, $p$ = 0.20) or left ventricular thickness (one-way ANOVA; $F_{2,12}$ = 1.25, $p$ = 0.32) between genotypes. Scale bar for **A** is 500 µm, and the scale bars for **B-C** are 50 µm. Data is presented as mean ± SEM.

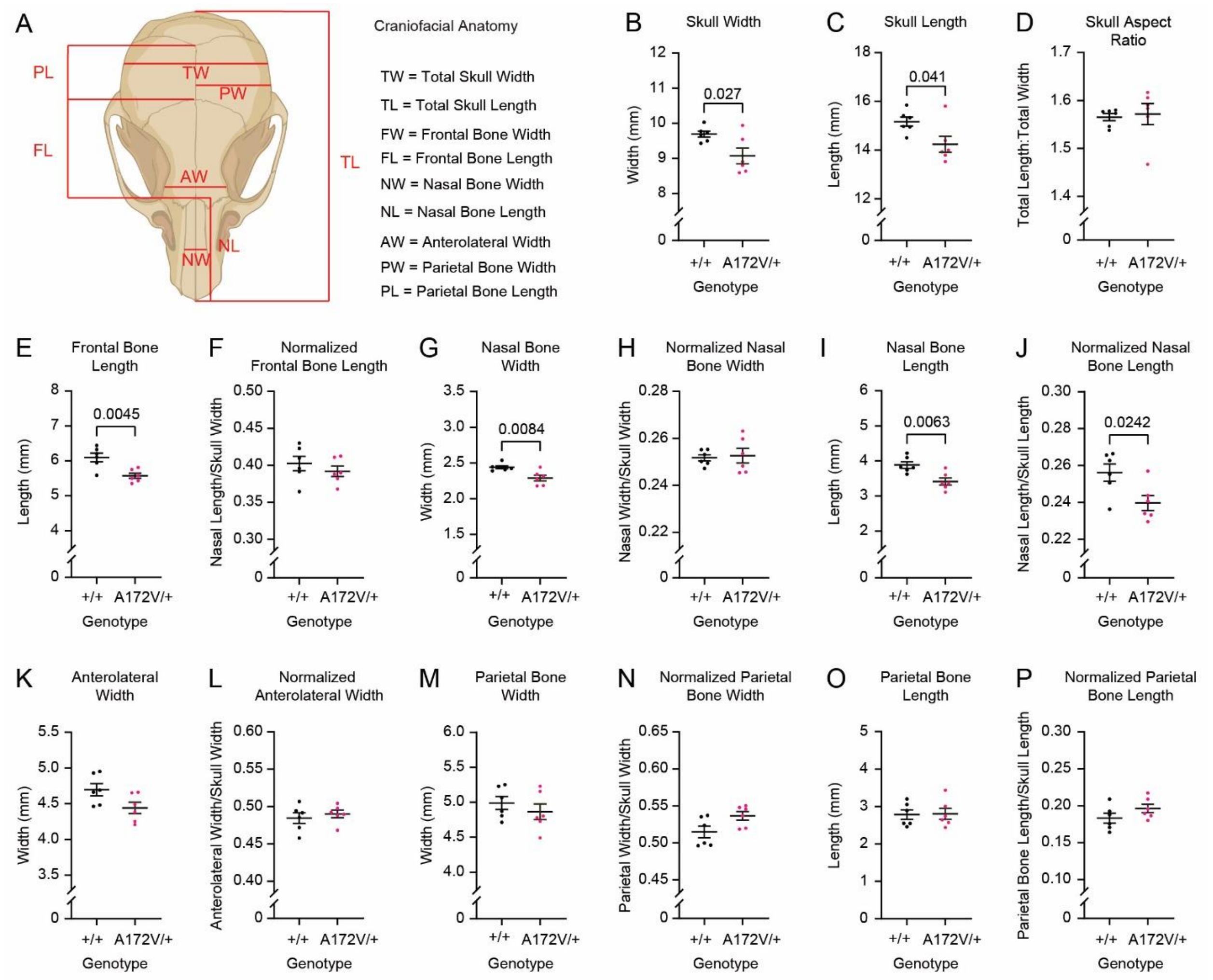


**Supplementary Figure 4. Craniofacial analysis in *Mapk1* mutant and control mice. (A)** Schematic showing skull features measures in PD7 *Mapk1*$^{+/+}$ and *Mapk1*$^{A172V/+}$ mice. **(B-C)** There was a significant reduction in the overall width ($n_{+/+}$ = 6; $n_{A172V/+}$ = 6; unpaired t-test; $t_{10}$= 2.59, $p$ = 0.027) and length of skulls in heterozygous mutant mice (Mann-Whitney U-test; $U(n_{+/+} = 6, n_{A172V/+} = 6) = 5$, $p$ = 0.041). **(D)** There was no difference in the skull aspect ratio (length:width) between genotypes (Mann-Whitney U-test; $U(n_{+/+} = 6, n_{A172V/+} = 6) = 10$, $p$ = 0.24). **(E-F)** There was a significant reduction in frontal bone length in *Mapk1*$^{A172V/+}$ mice ($n_{+/+}$ = 6; $n_{A172V/+}$ = 6; unpaired t-test; $t_{10}$= 3.65, $p$ = 0.0045) that was not significant after normalizing to the total skull length ($n_{+/+}$ = 6; $n_{A172V/+}$ = 6; unpaired t-test; $t_{10}$= 0.88, $p$ = 0.40). **(G-H)** There was a significant reduction in nasal bone width ($n_{+/+}$ = 6; $n_{A172V/+}$ = 6; unpaired t-test; $t_{10}$= 3.27, $p$ = 0.0084) in *Mapk1*$^{A172V/+}$ mice that was not significant after normalizing to the overall skull width ($n_{+/+}$ = 6; $n_{A172V/+}$ = 6; unpaired t-test; $t_{10}$= 0.27, $p$ = 0.80). **(I-J)** There was a significant decrease in the nasal bone length in *Mapk1*$^{A172V/+}$ mice compared to wildtype littermates ($n_{+/+}$ = 6; $n_{A172V/+}$ = 6; unpaired t-test; $t_{10}$= 3.44, $p$ = 0.0063) that persisted after it was normalized to the total skull length ($n_{+/+}$ = 6; $n_{A172V/+}$ = 6; unpaired t-test; $t_{10}$= 2.65, $p$ = 0.024). **(K-P)** There were no significant differences in the skull anterolateral width ($n_{+/+}$ = 6; $n_{A172V/+}$ = 6; unpaired t-test; $t_{10}$= 2.19, $p$ = 0.054), normalized anterolateral width ($n_{+/+}$ = 6; $n_{A172V/+}$ = 6; unpaired t-test; $t_{10}$= 0.63, $p$ = 0.54), parietal bone width ($n_{+/+}$ = 6; $n_{A172V/+}$ = 6; unpaired t-test; $t_{10}$= 0.86, $p$ = 0.41), normalized parietal bone width (Mann-Whitney U-test; $U(n_{+/+}$ = 6, $n_{A172V/+}$ = 6) = 7, $p$ = 0.093), parietal bone length (unpaired t-test; $t_{10}$= 0.11, $p$ = 0.92), and normalized parietal bone length ($n_{+/+}$ = 6; $n_{A172V/+}$ = 6; unpaired t-test; $t_{10}$= 1.48, $p$ = 0.17) between *Mapk1*$^{A172V/+}$ and *Mapk1*$^{+/+}$ mice. Data is presented as mean ± SEM.

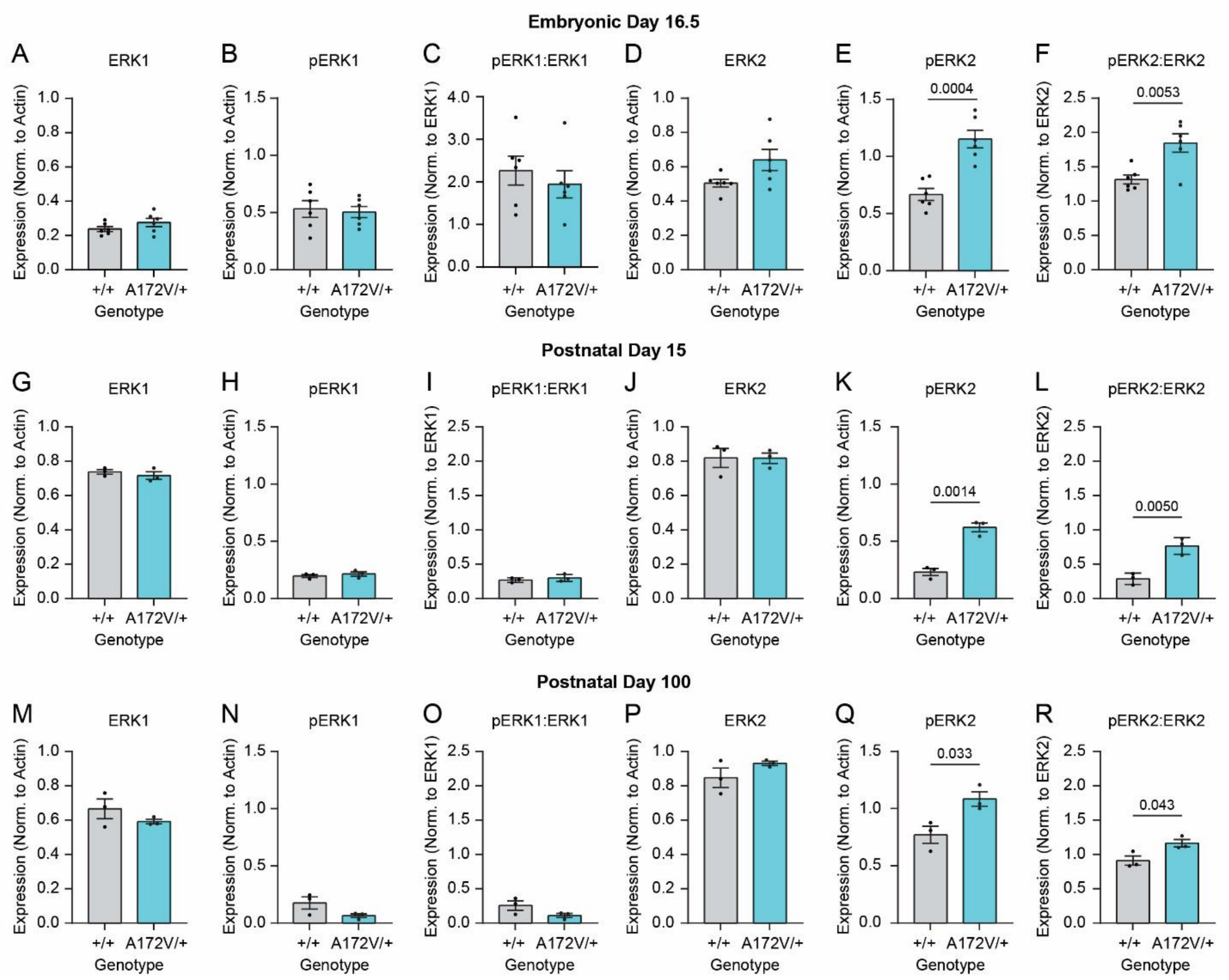


**Supplementary Figure 5. Developmental time course of ERK phosphorylation in the brains of *Mapk1* mutant and control mice. (A-D)** At ED16.5, there were no genotypic differences in the level of ERK1 ($n_{+/+}$ = 6; $n_{A172V/+}$ = 6; unpaired t-test; $t_{10}$ = 1.33, $p$ = 0.21), pERK1 ($n_{+/+}$ = 6; $n_{A172V/+}$ = 6; unpaired t-test; $t_{10}$ = 0.305, $p$ = 0.77), the pERK1:ERK1 ratio ($n_{+/+}$ = 6; $n_{A172V/+}$ = 6; unpaired t-test; $t_{10}$ = 0.69, $p$ = 0.51), or total ERK2 ($n_{+/+}$ = 6; $n_{A172V/+}$ = 6; unpaired t-test with Welch's correction; $t_{10}$ = 2.073, $p$ = 0.082) in forebrain samples, as measured by Western blot. **(E-F)** There were significant increases in pERK2 ($n_{+/+}$ = 6; $n_{A172V/+}$ = 6; unpaired t-test; $t_{10}$ = 5.18, $p$ = 0.0004) and the pERK2:ERK2 ratio ($n_{+/+}$ = 6; $n_{A172V/+}$ = 6; unpaired t-test; $t_{10}$ = 3.55, $p$ = 0.0053) in *Mapk1*$^{A172V/+}$ forebrain samples relative to controls at ED16.5. **(G-J)** At PD15, there were no significant genotypic differences in the abundance of ERK1 ($n_{+/+}$ = 3; $n_{A172V/+}$ = 3; unpaired t-test;

$t_4 = 0.83$, $p = 0.45$), pERK1 ($n_{+/+} = 3$; $n_{A172V/+} = 3$; unpaired t-test; $t_4 = 0.72$, $p = 0.51$), pERK1:ERK1 ($n_{+/+} = 3$; $n_{A172V/+} = 3$; unpaired t-test; $t_4 = 0.89$, $p = 0.42$), and ERK2 ($n_{+/+} = 3$; $n_{A172V/+} = 3$; unpaired t-test; $t_4 = 0.025$, $p = 0.98$) in forebrain samples. **(K-L)** Significant increases in pERK2 ($n_{+/+} = 3$; $n_{A172V/+} = 3$; unpaired t-test; $t_4 = 7.88$, $p = 0.0014$) and pERK2:ERK2 ($n_{+/+} = 3$; $n_{A172V/+} = 3$; unpaired t-test; $t_4 = 5.58$, $p = 0.005$) were observed in *Mapk1*$^{A172V/+}$ forebrain samples at PD15. **(M-P)** At PD100, there were no genotypic differences in ERK1 ($n_{+/+} = 3$; $n_{A172V/+} = 3$; unpaired t-test; $t_4 = 1.26$, $p = 0.28$), pERK1 ($n_{+/+} = 3$; $n_{A172V/+} = 3$; unpaired t-test; $t_4 = 1.99$, $p = 0.12$), pERK1:ERK1 ($n_{+/+} = 3$; $n_{A172V/+} = 3$; unpaired t-test; $t_4 = 1.93$, $p = 0.13$), or ERK2 ($n_{+/+} = 3$; $n_{A172V/+} = 3$; unpaired t-test; $t_4 = 1.47$, $p = 0.22$) in forebrain samples. Significant increases in pERK2 ($n_{+/+} = 3$; $n_{A172V/+} = 3$; unpaired t-test; $t_4 = 3.204$, $p = 0.033$) and pERK2:ERK2 ($n_{+/+} = 3$; $n_{A172V/+} = 3$; unpaired t-test; $t_4 = 2.93$, $p = 0.043$) were observed in *Mapk1*$^{A172V/+}$ forebrain samples at PD100. Data is presented as mean ± SEM.

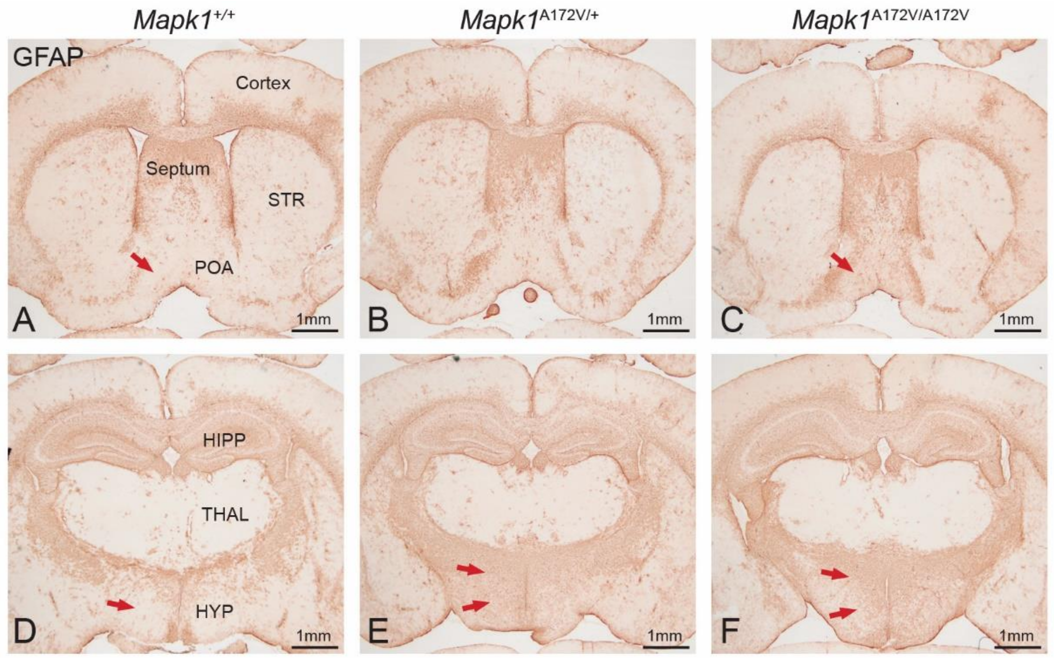


**Supplementary Figure 6. GFAP expression in the brains of *Mapk1* mutant and control mice. (A-C)** Images showing GFAP expression (a marker for reactive astrocytes) in different brain structures in 10-week-old **(A)** *Mapk1*$^{+/+}$, **(B)** *Mapk1*$^{A172V/+}$, and **(C)** *Mapk1*$^{A172V/A172V}$ mice, including the cerebral cortex, septum, striatum (STR), and pre-optic area (POA). Red arrows indicate differences in GFAP immunoreactivity in the pre-optic area. **(D-F)** Images showing GFAP expression in **(D)** *Mapk1*$^{+/+}$, **(E)** *Mapk1*$^{A172V/+}$, and **(F)** *Mapk1*$^{A172V/A172V}$ mice in the hippocampus (HIPP), thalamus (THAL), and hypothalamus (HYP). Red arrows indicate differences in GFAP immunoreactivity in the hypothalamus. Scale bars are 1mm.

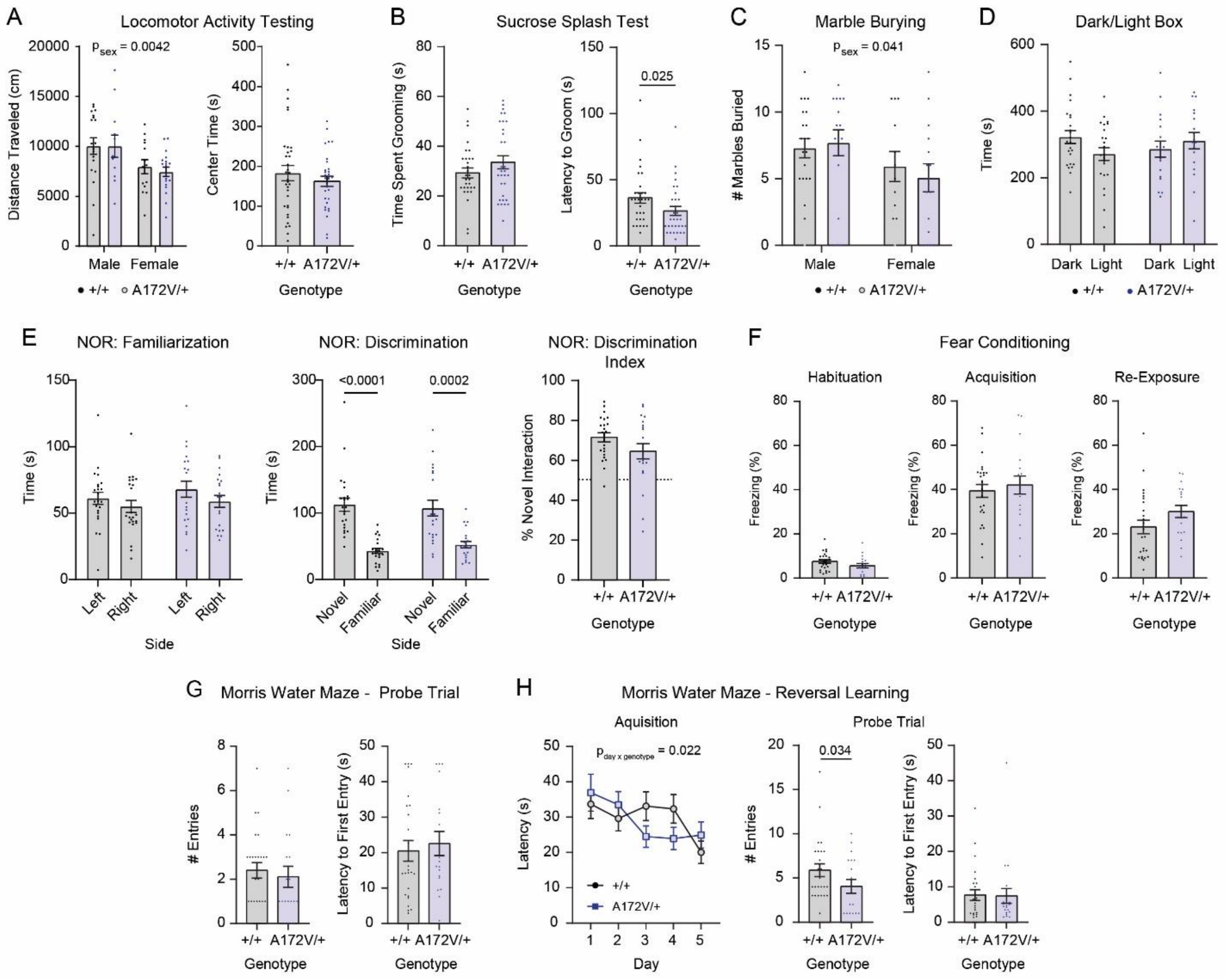


**Supplementary Figure 7. Behavioral testing of *Mapk1* mutant and control mice. (A)** In a test of locomotor activity, there was a main effect of sex but not genotype (*left*; $n_{+/+}$ = 33, $n_{A172V/+}$ = 31; two-way ANOVA; $F_{1,60}$ = 0.098, $p_{sex\ x\ genotype}$ = 0.76; $F_{1,60}$ = 8.87, $p_{sex}$ = 0.0042; $F_{1,60}$ = 0.12, $p_{genotype}$ = 0.73), and there were no changes in the time spent in the center of the arena (*right*; $n_{+/+}$ = 33, $n_{A172V/+}$ = 31; unpaired t-test with Welch's correction; $t_{55.23}$= 0.89, $p$ = 0.38). **(B)** *Mapk1*$^{A172V/+}$ mice showed no significant changes in the amount of time spent grooming in the sucrose splash test (*left*; Mann-Whitney U-test; $U(n_{+/+}$ = 30, $n_{A172V/+}$ = 31) = 408, $p$ = 0.42) but had a shorter latency to initiate grooming behavior (*right*; Mann-Whitney U-test; $U(n_{+/+}$ = 30, $n_{A172V/+}$ = 31) = 311.5, $p$ = 0.025). **(C)** There was a significant main effect of sex but not genotype in the marble burying assay ($n_{+/+}$ = 33; $n_{A172V/+}$ = 31; two-way ANOVA; $F_{1,60}$ = 0.43, $p_{sex\ x\ genotype}$ = 0.52; $F_{1,60}$ = 4.35, $p_{sex}$

= 0.041; $F_{1,60}$ = 0.055, $p_{genotype}$ = 0.82). **(D)** There were no effects of genotype or side in the dark/light box assay ($n_{+/+}$ = 24; $n_{A172V/+}$ = 18; two-way ANOVA; $F_{1,80}$ = 3.075, $p_{side\ x\ genotype}$ = 0.083; $F_{1,80}$ = 0.35, $p_{side}$ = 0.56; $F_{1,80}$ = 0.0072, $p_{genotype}$ = 0.93). **(E)** No genotypic differences in object preference were observed during the familiarization phase of the novel object recognition (NOR) test (*left*; $n_{+/+}$ = 23; $n_{A172V/+}$ = 21; two-way repeated measures ANOVA; $F_{1,42}$ = 0.12, $p_{side\ x\ genotype}$ = 0.74; $F_{1,42}$ = 2.94, $p_{side}$ = 0.094; $F_{1,42}$ = 1.021, $p_{genotype}$ = 0.32). During the discrimination phase, both genotypes interacted more with the novel object (*middle*; $n_{+/+}$ = 23; $n_{A172V/+}$ = 21; two-way ANOVA; $F_{1,42}$ = 0.69, $p_{object\ x\ genotype}$ = 0.41; $F_{1,42}$ = 49.37, $p_{object}$ < 0.0001; $F_{1,42}$ = 0.065, $p_{genotype}$ = 0.80). There were no differences in the object discrimination index ((time spent with novel object – time spent with familiar objects) / total time interacting with both objects) between genotypes (*right*; $n_{+/+}$ = 23; $n_{A172V/+}$ = 21; unpaired t-test with Welch's correction; $t_{33.03}$= 1.58, $p$ = 0.12). **(F)** During contextual fear conditioning, there were no significant differences between genotypes during the habituation phase before shocks were delivered (*left*; $n_{+/+}$ = 25, $n_{A172V/+}$ = 18; unpaired t-test; $t_{41}$= 1.55, $p$ = 0.13), acquisition phase when shocks were presented (*middle*; $n_{+/+}$ = 25, $n_{A172V/+}$ = 18; unpaired t-test; $t_{41}$= 0.54, $p$ = 0.59), or during context re-exposure 24-hours later (r*ight*; $n_{+/+}$ = 25, $n_{A172V/+}$ = 18; unpaired t-test; $t_{41}$= 1.62, $p$ = 0.11). **(G)** During the probe trial of the Morris water maze (MWM), there were no significant differences in the number of entries into the target quadrant (*left*; Mann-Whitney U-test; $U(n_{+/+}$ = 25, $n_{A172V/+}$ = 18) = 194.5, $p$ = 0.42) or latency to the first entry into the target quadrant (*right*; Mann-Whitney U-test; $U(n_{+/+}$ = 25, $n_{A172V/+}$ = 18) = 199, $p$ = 0.53). **(H)** During the reversal learning acquisition phase of the MWM, there was a significant interaction between day and genotype (left; $n_{+/+}$ = 25, $n_{A172V/+}$ = 17; two-way ANOVA; $F_{4,160}$ = 2.95, $p_{day\ x\ genotype}$ = 0.022; $F_{3.73,149.1}$ = 5.26, $p_{day}$ = 0.0007; $F_{1,40}$ = 0.024, $p_{genotype}$ = 0.88). *Post hoc* analysis determined that in wild-type mice, there were significant differences in latency on day 5 compared to their latency on days 1, 3, and 4 (Bonferroni *post hoc* test; $p$ < 0.05), while no differences were observed in *Mapk1*$^{A172V/+}$ mice (Bonferroni *post hoc* test, $p > 0.05$). During the reversal learning probe trial, there were no differences in entries into the target quadrant (*middle*; Mann-Whitney U-test; $U(n_{+/+} = 25, n_{A172V/+} = 15) = 120.5$, $p = 0.060$) and no difference in the latency to enter the target quadrant (right; Mann-Whitney U-test; $U(n_{+/+} = 25, n_{A172V/+} = 15) = 184$, $p = 0.93$). Data is presented as mean ± SEM.
